# KH-type splicing regulatory protein controls colorectal cancer cell growth and modulates the tumor microenvironment

**DOI:** 10.1101/342840

**Authors:** Francesco Caiazza, Katarzyna Oficjalska, Miriam Tosetto, James J. Phelan, Sinéad Noonan, Petra Martin, Kate Killick, Laura Breen, Fiona O’Neill, Blathnaid Nolan, Simon Furney, Robert Power, David Fennelly, Charles S. Craik, Jacintha O’Sullivan, Kieran Sheahan, Glen A. Doherty, Elizabeth J. Ryan

**Author notes:** Corresponding author: Francesco Caiazza, Department of Pharmaceutical Chemistry, University of California San Francisco, 600 16^th^ street, Room S-514, San Francisco, California, U.S.A. Dept. of Biological Sciences, School of Engineering and Science, University of Limerick, Limerick, Ireland (Elizabeth J. Ryan). **Disclosures:** None declared. **Abbreviations:** KHSRP, KH-type splicing regulatory protein; CRC, colorectal cancer; IHC, immunohistochemistry; FFPE, formalin fixed paraffin embedded; RBP, RNA-binding protein; UTR, untranslated region; ARE, AU-rich element; TCGA, The cancer genome atlas; DAMP, damage-associated molecular pattern; PAMP, pathogen-associated molecular pattern.

## Abstract

KH-type splicing regulatory protein (KHSRP) is a multifunctional nucleic acid binding protein implicated in key aspects of cancer cell biology: inflammation and cell-fate determination. However, the role KHSRP plays in colorectal cancer (CRC) tumorigenesis remains largely unknown. Using a combination of *in silico* analysis of large datasets, *ex vivo* analysis of protein expression in patients, and mechanistic studies using *in vitro* models of CRC, we investigated the oncogenic role of KHSRP. We demonstrated KHSRP expression in the epithelial and stromal compartments of both primary and metastatic tumors. Elevated expression was found in tumor versus matched normal tissue, and we validated these findings in larger independent cohorts *in silico.* KHSRP expression was a prognostic indicator of worse overall survival (HR=3.74, 95% CI = 1.43-22.97, p=0.0138). Mechanistic data in CRC cell line models supported a role of KHSRP in driving epithelial cell proliferation in both a primary and metastatic setting, through control of the G1/S transition. Additionally, KHSRP promoted a pro-angiogenic extracellular environment by regulating the secretion of oncogenic proteins involved in diverse cellular processes such as migration and response to cellular stress. Our study provides novel mechanistic insight into the tumor-promoting effects of KHSRP in CRC.

## INTRODUCTION

Almost all the major oncogenic signaling pathways result in the reprogramming of translation. This impacts on both cancer initiation and progression by directing selective translation of specific tumor-promoting mRNAs ^1^. RNA-binding proteins (RBP) such as KH-type splicing regulatory protein (KHSRP, also known as KSRP or FUBP2) are capable of recognizing sequence-specific regulatory elements in the 3’UTR, such as AU-rich elements (ARE) that regulate transcript specific translation promoting mRNA splicing and stability ^2^. KHSRP regulates translation by mediating mRNA decay through direct binding and recruitment of proteins involved in RNA degradation, including the Poly(A)-Specific Ribonuclease (PARN), exosome components and decapping enzymes ^3, 4^. Alternatively, KHSRP can regulate translation independently of mRNA decay, either by promoting translational inhibition ^5^ or indirectly by promoting maturation of selected miRNA precursors ^6^. Adding a further layer of complexity, KHSRP also acts as a transcription factor, inducing the expression of the *MYC* oncogene ^7^.

The KHSRP-mediated control of mRNA translation and stability has been extensively studied in the context of innate immunity ^5, 8^. Pro-inflammatory cytokines (e.g. IL-6, IL-8, TNF-α, IL-1β) and inflammatory mediators such as iNOS are directly targeted and negatively regulated by KHSRP ^9,10^. However, the role of KHSRP is most likely context dependent, with the balance between KHSRP and other RBP with similar or divergent regulatory effects an important consideration ^11^.

Direct evidence for the involvement of KHSRP in cancer is accumulating. KHSRP has been implicated in the pathogenesis of small cell lung cancer ^12^, osteosarcoma ^13^ and breast cancer ^14^. A number of distinct mechanisms have been proposed e.g. regulation of cell differentiation in P19 mouse teratocarcinoma cells ^15^, deregulation of oncosuppressive miRNAs such as let-7a and miR-30c ^16^ or control of TGF-β-dependent regulation of epithelial-to-mesenchymal transition ^17^.

The role of KHSRP in colorectal cancer (CRC), however, remains underappreciated. While other RBPs have been implicated in cellular transformation in the intestinal epithelium [namely, ESRP1 ^18^, Apobec-1 ^19^, Musashi (MSI)-1 ^20^, and MSI-2 ^21^], KHSRP remains largely under-investigated in this indication. Interestingly, adenomatous polyposis coli (APC), a tumor suppressor frequently mutated in CRC, is itself a RBP capable of regulating the translation of mRNAs associated with cell adhesion, motility and other cellular processes crucial for carcinogenesis ^22^, suggesting the importance of RBP-mediated translation control in the gut. Here we use a combination of bioinformatic, *in vitro* and *ex vivo* approaches to dissect the role of KHSRP in both regulation of cell proliferation and inflammatory environment in CRC and provide evidence for a novel prognostic role of this RBP in intestinal tumorigenesis.

## MATERIALS AND METHODS

### Bioinformatic analysis of publicly available datasets

CRC datasets in the Oncomine database (www.oncomine.org, accessed 11/2014, Thermo Fisher Scientific, Ann Arbor, MI) were searched using the differential expression module for cancer vs. normal tissue; output of the resulting database searches was downloaded for offline plotting, along with detailed gene expression results for 2 representative cohorts (GSE9348 and TCGA). A similar analysis was conducted without restricting for cancer type. Differential expression analysis of RNA-Seq data for 12 tumor-normal pairs in the TCGA database was previously described and made available by the authors ^23^. Gene expression (RNA-Seq) and the frequency of genetic alterations for KHSRP in the TCGA cohort were analyzed using the cBio portal (www.cbioportal.org, accessed 11/2016) ^24^. All other datasets were downloaded from the Gene Expression Omnibus (https://www.ncbi.nlm.nih.gov/geo/), using the indicated accession number.

### Patient Samples and Tissue Microarrays

Fresh-frozen tumor and adjacent normal tissue samples were obtained from 16 patients and were used for protein isolation and Western blotting (WB cohort) as described thereafter. Tissue microarrays (TMA) were accessed from a previously described cohort of formalin-fixed paraffin-embedded (FFPE) archival resected tumor tissue samples ^25^. Patient details for both cohorts are reported in Table 1. For immunohistochemistry, TMA sections were incubated with a 1:250 dilution of a validated (https://www.proteinatlas.org/ENSG00000088247-KHSRP/antibody, accessed 04/2019) rabbit polyclonal anti-KHSRP antibody (Cat# HPA034739, RRID:AB_10601582, Sigma-Aldrich, Saint Louis, MO, USA), and secondary antibody horseradish peroxidase (DAKO Agilent, Santa Clara, CA, USA). Endogenous peroxidase activity was blocked using 3% hydrogen peroxide, and non-specific binding was blocked with casein buffer. Diaminobenzidine was used to visualize staining and sections were counter-stained with haematoxylin, dehydrated and mounted. Stained TMA sections were acquired with a ScanScope XT high throughput scanning system (Aperio Technologies, Leica Biosystems, Buffalo Grove, IL, USA) and images were scored by two independent investigators, blinded to the clinical data. Both epithelial and stromal cells were assessed for staining intensity and percent positivity; positive cell count was scored as a continuous variable (0-100%), while intensity was scored using a scale of 0 to 3 (correlating with negative, weak, medium and strong staining, respectively). The mean value of each parameter from the two investigators was calculated, and parameters were averaged from all tissue cores related to the same patient; the two parameters were finally combined using the semi-quantitative *quickscore* method ^26^, by multiplying the averaged intensity and positivity scores for epithelial and stromal cells separately. The quickscore values for tumor tissue were further divided by the quickscore values for normal tissue, to obtain a tumor-to-normal (T/N) ratio score that was used to quantify the amplitude of the tumor-specific changes in protein expression. A similar strategy was used to obtain metastasis-to-normal (M/N) scores. There was overall a good correlation between the scores obtained by the two independent assessors (weighted Cohen Kappa coefficient = 0.63).

**Table 1:**
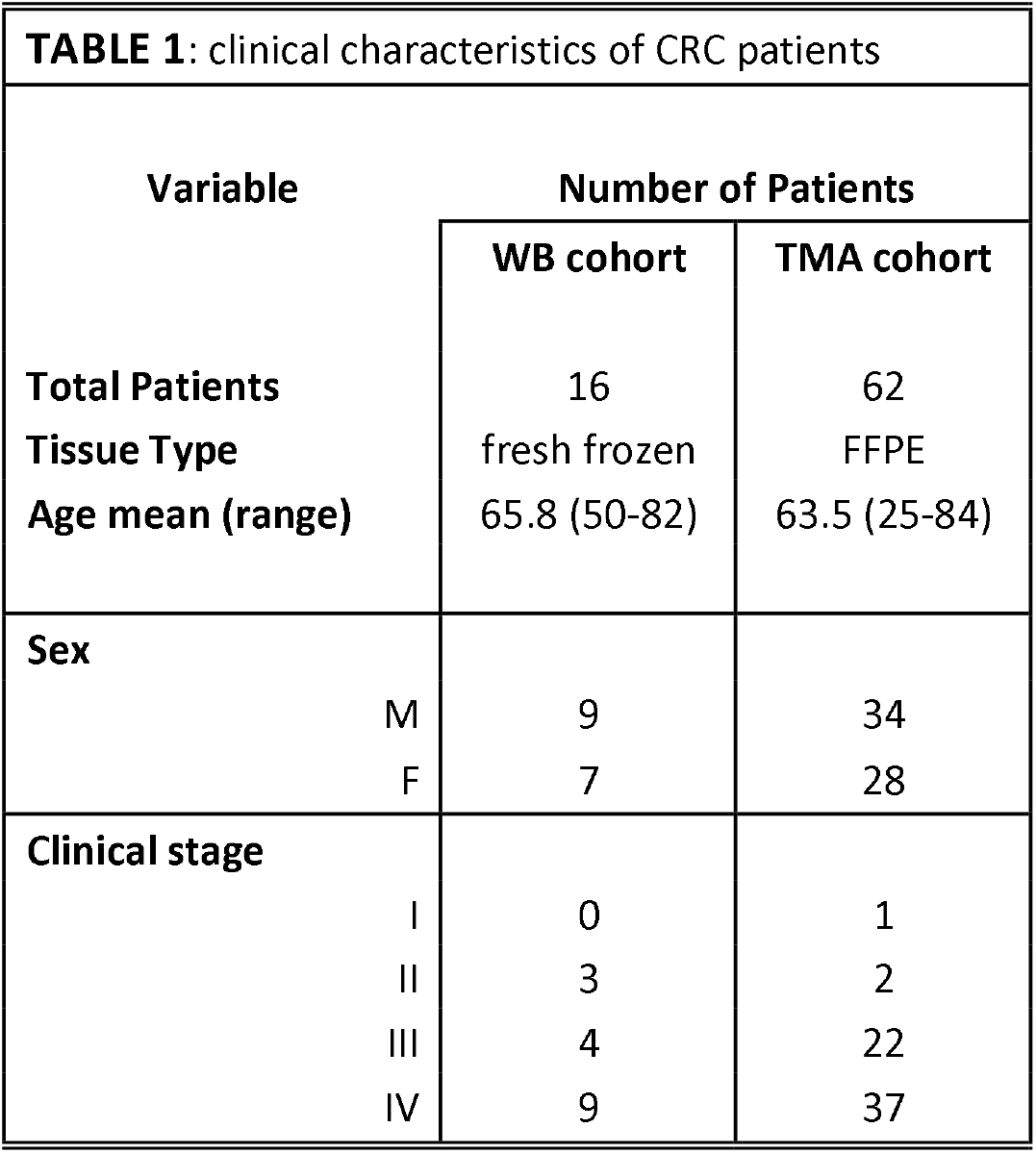
Summary of clinical characteristics for the patients included in the two cohorts analyzed by either Western blotting (WB) or Tissue Microarrays (TMA).

### Cell Culture, transient and stable knockdown of KHSRP

SW480 and SW620 cells were obtained from the American Tissue Culture Collection (ATCC, Manassas, VA, USA) and maintained in DMEM supplemented with 10% fetal bovine serum, 1% penicillin/streptomycin and 1% Fungizone (all from Invitrogen Life Technologies, Carlsbad, CA, USA). Cell line identity was confirmed by analysis of Short Term Repeat loci, and cells were routinely tested for mycoplasma infection. Cells were seeded in 24-well plates to be transfected with 30 pmol KHSRP siRNA (Silencer Select, Thermo Fisher) or non-targeting scramble siRNA using 2 μl Lipofectamine 2000 (Invitrogen) in serum-free and antibiotic-free growth media for 6h, followed by incubation in standard growth media for a total of 48h post-transfection. Alternatively, cells were transfected with 50 nM KHSRP siRNA (SMARTpool: ON-TARGET plus) and corresponding Non-Targeting Pool (Dharmacon, Lafayette, CO, USA). To generate stable KHSRP knockdown cell lines, we took advantage of an ultracomplex pooled shRNA library targeting each annotated human protein-coding gene with 25 independent shRNAs on average (as well as 500 negative control shRNAs), which was previously described and used to perform genome-wide genetic interaction screens in mammalian cells ^27^. From this original library, 3 independently validated shRNA sequences targeting KHSRP were selected, along with one negative control scramble shRNA sequence; for each sequence, top and bottom oligonucleotides were synthesized (Integrated DNA Technologies, Coralville, Iowa, USA). Two shRNA-expressing derivative cell lines were generated from SW480 and SW620 following a previously published protocol ^28^. The lentiviral plasmid used for shRNA expression was pMK1201, which is a modified version of pMK1200 ^28^ based on the design of pINDUCER10 for tetracycline-induced expression of the shRNA coupled with a fluorescent reporter (tRFP) and a Puromycin resistance sequence for positive selection ^29^. To clone each individual shRNA into the lentiviral backbone, 2 μl of top and bottom oligonucleotides were annealed at 95°C for 5 min in a buffer containing 100 mM potassium acetate, 30 mM HEPES-KOH (pH=7.4), 2 mM magnesium acetate. Annealed oligonucleotides (0.01 μM) were ligated in 25 ng of the pMK1201 vector pre-digested with BstXI (New England Biolabs, Ipswich, MA, USA) and gel purified; ligation was carried out at RT for 2h using 2000 U of T4 DNA ligase (New England Biolabs). DH5α cells (Thermo Fisher) were transformed and plated onto ampicillin-containing LB plates overnight at 37°C. Single colonies were picked and expanded for plasmid DNA mini prep (Qiagen, Germantown, MA, USA). Correct insertion of shRNA sequences was confirmed by sequencing using the 5’ pSico-Eco-insert-seq ^28^. For lentivirus preparation, the second-generation virion packaging vector psPAX2 (plasmid #12260, Addgene, Cambridge, MA, USA) and the VSV-G envelope plasmid pMD2.G (plasmid #12259, Addgene) from Didier Trono (Ecole Polytechnique Fédérale de Lausanne) were used. The producer cell line was 293T, a highly-transfectable derivative of human embryonic kidney cell 293 (ATCC), which was maintained in DMEM supplemented with 10% fetal bovine serum, 1% penicillin/streptomycin and 1% Fungizone (Invitrogen). 293T cells were transfected in 10 cm plates using the calcium phosphate method: 2.5 μg of scramble shRNA vector, or a pooled mix of 0.83 μg each of the three KHSRP shRNA vectors, were combined with 0.58 μg of pMD2.G and 1.92 μg of psPAX2 and transfected with 12.5 μM chloroquine diphosphate and 12.5 mM of calcium chloride. Transfection medium was replaced after 16h with full DMEM, and virus-containing conditioned medium was harvested 2 times at 48h and 72h post transfection; pooled harvests from each condition were pre-cleared by centrifugation, filtered through a 0.45 μm membrane, and used immediately to transduce SW480 and SW620. Cells were selected with 1 μg/ml puromycin for 2 weeks, expanded and stock-frozen. A fresh stock of cells was used each time for experiments.

### Protein isolation, immunoblotting, immunofluorescence and ELISA

Cells or tissue homogenate were lysed in RIPA buffer (150 mM NaCl, 50 mM Tris-HCl, 1% Triton, 0.5% sodium deoxycholate, 0.1% SDS) supplemented with a protease and phosphatase inhibitor cocktail (Roche Applied Science, Burgess Hill, UK) and 1 mM PMSF (Sigma Aldrich). Total protein concentration was determined using the BCA Protein Assay Kit (Thermo Fisher). Total proteins (10-20μg) were separated on 10% or 4-20% gradient SDS–PAGE gels and wet-transferred to PVDF using the mini-PROTEAN^®^ TGX^™^ system (Bio-Rad, Hercules, CA, USA). Membranes were pre-blocked with 5% low-fat dry milk in TBS-T and incubated with a rabbit polyclonal anti-KHSRP antibody (Sigma-Aldrich Cat# HPA034739, RRID:AB_10601582), or a monoclonal anti-c-Myc antibody (D84C12, Cell Signaling, Danvers, MA, USA), or a monoclonal anti-EphB2 (clone D2X2I, Cell Signaling), or a monoclonal anti-S100A11 (clone EPR11171B, Abcam, Cambridge, MA, USA), and either anti-rabbit or anti-mouse horseradish peroxidase-conjugated secondary antibody (Bio-Rad). Protein loading normalization was carried out with anti-β-actin or anti-GAPDH antibodies (Cell Signaling), or Coomassie stain (Thermo Fisher). Proteins were visualized by chemiluminescence with ECL substrate (Thermo Fisher) and semi-quantified using ImageJ software (US National Institute of Health, Bethesda, MD, USA; http://imagej.nih.gov/ij/, version 2.0.0-rc-69/1.52n) with normalization against β-actin (Sigma-Aldrich). For immunofluorescence, cells were fixed on chamber slides with 4% Formaldehyde, permeabilized with 0.5% Tween in PBS, and stained using the same anti-KHSRP antibody and a Cy3-anti-rabbit, with DAPI and Phalloidin counter-stain (DAKO); slides were imaged on a Leitz DM40 microscope (Leica Microsystems) equipped with the AxioCam system and AxioVision 3.0.6 for image acquisition. Human interleukin (IL)-8 and vascular endothelial growth factor (VEGF) were quantified in cell culture supernatants using DuoSet^®^ ELISA (R&D Systems, Minneapolis, MN, USA), following manufacturer’s instructions.

### RNA isolation and qRT-PCR

Total RNA was isolated from cells using the EZNA Total RNA Kit (Omega, VWR, Radnore, PA, USA). 1 μg of RNA was reverse-transcribed into cDNA using the RT Omniscript cDNA kit (Qiagen), and 20 ng of cDNA template was amplified on a LightCycler-480 (Roche, Indianapolis, IL, USA) using a FAST SYBR Green Master Mix (Invitrogen) under the following conditions: pre-incubation step at 95 °C for 5 min was followed by 45 cycles of denaturation at 95 °C for 10 sec, annealing at 60 °C for 10 sec, and elongation at 72 °C for 10 sec. GAPDH was used as the housekeeping gene. The mRNA expression levels of all samples were normalised to the housekeeping gene, and the delta/delta Ct method was used to calculate relative expression (treatment vs. control). The following primers were used in qPCR: KHSRP Forward: 5’-CCGCTTACTACGGACAGACC-3’; KHSRP Reverse: 5’-ATTCATTGAGCCTGCTGCTGT-3’; GAPDH Forward: 5’-CGGATTTGGTCGTATTGGGCGCCTG-3’; GAPDH Reverse: 5’-CAGCATCGCCCCACTTGATTTTGGA-3’. For validation of microarray candidate genes, single strand cDNA synthesis and PCR amplification were carried out in a 1-step reaction using the Brilliant II QRT-PCR Master Mix (Agilent) and TaqMan gene expression assays (Life Technologies). 25 ng of RNA was loaded in 96-well plate format and amplified with 0.9 mM of primers and 0.25 mM of MGB probe. The reaction was carried out in a CFX Connect instrument (Bio-Rad) with the following parameters: an RT step at 50 LJC for 30 min was followed by a preincubation step at 95 LJC for 10 min, 50 cycles of denaturation at 95 LJC for 15 s, and annealing/extension at 60 LJC for 1 min. Relative quantification of gene expression was calculated with the comparative cycle threshold method, normalizing for GAPDH expression levels. TaqMan gene expression assays for FABP3 (GeneID: 2170, Assay ID: Hs00997360_m1), SMG1 (GeneID: 23049, Assay ID: Hs00979691_m1), MT1F (GeneID: 4494, Assay ID: Hs00744661_sH), and GAPDH (GeneID: 2597, Assay ID: Hs02786624_g1) contained MGB probes with a FAM reporter dye.

### Gene Expression Microarray

Total RNA was extracted from cells using the RNeasy Plus kit (Qiagen); the integrity and concentration of RNA was confirmed using the RNA 6000 Nano kit on a Bioanalyzer 2100 (Agilent), with reported RNA Integrity Numbers (RIN) > 9. Amplified cDNA for gene expression analysis was prepared with the Ovation® PicoSL WTA System V2 (Nugen, San Carlos, CA, USA); labeled cDNA targets were generated with the Encore® Biotin Module (Nugen) and hybridized to a GeneChip® Human Transcriptome Array 2.0 (Affymetrix, Thermo Fisher) following manufacturer’s instructions. CEL files were analysed with the Affymetrix® Expression Console™ Software, using the Affymetrix Human Transcripome Array 2.0 library files and the HTA-2_0.na35.2.hg19 annotation files. An RMA workflow was performed which used a quantile normalization procedure and a general background correction. The resulting normalized CHP files were then imported in the Affymetrix® Transcriptome Analysis Console software to test for differential expression using a paired One-Way Repeated Measure (ANOVA) approach, and the default filter criteria (fold change > |2|, and p-value < 0.05). The full data have been deposited in NCBI’s Gene Expression Omnibus, and can be accessed with the GEO Series accession number GSE112329. Over-representation analysis was used to identify canonical pathways and functional processes of biological importance within the list of differentially expressed genes; the analysis was performed using GeneTrail2 (version 1.5, https://genetrail2.bioinf.uni-sb.de, accessed 04/2016) ^30^. Network analysis was performed using String (version 10.0, https://string-db.org/, accessed 04/2016) ^31^. STRING is a database of known and predicted protein-protein interactions, including both direct (physical) and indirect (functional) associations, inferred from experimental data and computational predictions. STRING computes a global score by combining the probabilities from the different types of evidence and correcting for the probability of randomly observing an interaction. Proteins in the network are then clustered based on the distance matrix obtained from the global scores, using the KMEANS clustering algorithm ^32^.

### Cell-based assays

Cell proliferation was measured by Neutral Red (NR) staining following an established protocol ^33^. Briefly, cells were seeded in 24-well plates (20,000 cells/well) 24h after transfection (for the transient knockdown) or 24h after doxycycline treatment (for the stable knockdown), and allowed to grow for 7 days. Cells were stained with 33 mg/L neutral red (Sigma) for 2h, then washed in PBS and imaged (for representative photographs of the plates); cells were de-stained using a solution of 50% ethanol and 1% glacial acetic acid in water, and absorbance measured at 540nm to quantify staining.

Real-time cell proliferation was measured using an E-plate 16 and a RTCA DP Analyser (xCelligence System, Roche), which uses gold microelectrodes fused to the bottom surface of a microtiter plate well. The impedance of electron flow at the interface between electrodes and culture medium caused by adherent cells is reported using a unitless parameter called the Cell Index (CI). Cells were seeded at 10,000 cells/well and the recording of the CI occurred every 15 min during the first six hours, and every hour for the rest of the period. Cells seeded at low density (5,000 cells/well in 6-well plates) for colony-formation assay were grown for 7 days, then fixed in methanol/citric acid (4:1 v/v), stained in 2% crystal violet and counted with ImageJ. The plating efficiency was calculated by dividing the number of colonies by the original seeding density, and the surviving fraction was determined by comparing the plating efficiency of transfected vs control wells. The total colony area was calculated for each biological replicate by averaging the area of all colonies in replicate wells. Spheroids were generated by seeding 20,000 cells/well in 24-well plates on growth factor-reduced Matrigel (BD Biosciences, San Jose, CA, USA) following the 3D on-top protocol ^34^. After 7 days, spheroids were stained with neutral red (as outlined above), imaged (averaging 3-4 fields per well at 10X magnification) and quantified via ImageJ. Scratch-wound assays were performed in culture inserts 2-well μ-dishes (Ibidi, Martinsried, Germany), quantified with ImageJ (using MiToBo plugin) and representative images prepared with WimScratch software (Ibidi). Invasion assays were performed using Matrigel-coated Boyden chambers (8-μm pore size; BioCoat, BD Biosciences) for 72h against 10% FBS as chemo-attractant. Invaded cells were fixed using 1% glutaraldehyde, stained with 0.1% crystal violet, and imaged with a light microscope at a 10X magnification, averaging five random fields per insert. Crystal violet staining was quantified with ImageJ. For tube-formation assays, Human dermal microendothelial cells (HDEC, a gift from Prof. Ursula Fearon, TCD, Dublin, Ireland) were seeded on Matrigel-coated 96-well plates (10,000 cells/well) in MCDB131 medium (Invitrogen) with 40% conditioned medium from SW620 cells. The tube analysis was performed manually after 6h by counting five sequential fields (10X magnification) with a focus on the surface of the Matrigel, and a connecting branch between two discrete endothelial cells was counted as 1 tube. Representative images were prepared with WimTube (Ibidi).

### Flow Cytometry

Cell cycle analysis was carried out by propidium iodide (PI) staining. Cells were resuspended at 1-2×10^6^ cells/ml in PBS and fixed in 70% ethanol for 2h at −20°C, washed twice in PBS, and re-suspended in staining solution [0.1% (v/v) Triton X-100, 10 μg/mL PI, 100 μg/mL DNase-free RNase A in PBS] for 30’ at R.T. Data was acquired on a CyAn ADP (Beckman Coulter, Brea, CA, USA) using 488nm excitation laser, and data de-convolution was performed in FlowJo^®^ (Becton Dickinson, Franklin Lakes, NJ, USA). To quantify the mitotic index, fixed cells were permeabilized with 0.25% Triton X-100 in PBS for 20’ on ice, blocked with 10% BSA in PBS and incubated with 1μg of an Alexa-Fluor^®^ 488-conjugated anti-phospho-histone H3 (Ser10) antibody (Cat# 650804, BioLegend, San Diego, CA, USA). After washing, cells were re-suspended in PBS containing 50 μg/ml PI for 15’ on ice prior to analysis on a CyAn ADP using 488nm excitation laser. Mitotic cells with 4N DNA and elevated levels of phosphorylated histone H3 can be detected by plotting FL1-H versus FL3-A.

### Shotgun Proteomics

Stably transfected SW620 cells where treated with vehicle or 500ng/ml doxycycline for 4 days to induce shRNA-mediated knockdown of KHSRP, as outlined above. Cells were then incubated in serum-free medium for 16h, and triplicate conditioned media for each condition were harvested, buffer exchanged and concentrated in a final volume of 300 μl PBS using Amicon centrifugal filter units with a 3 kDa cut off membrane (MilliporeSigma, Burlington, MA, USA). From each sample, 10 μg total proteins were denatured with 6 M urea, disulfide bonds were reduced with 10 mM dithiothreitol (DTT), and free thiols were then alkylated with 12.5 mM iodoacetamide. The final volume was diluted 3-fold into 25 mM ammonium bicarbonate, and trypsin digestion was performed with a 1:50 mass ratio of sequencing-grade trypsin (Promega) to total protein for 16h at 37°C. Samples were acidified to approximately pH=2 with formic acid, and peptides desalted with C18 Desalting Tips (Rainin, Oakland, CA, USA), lyophilized, and rehydrated in 0.1% formic acid. Peptide sequencing by LC-MS/MS was performed on an LTQ Orbitrap Velos mass spectrometer (Thermo Fisher) equipped with a nanoACQUITY (Waters, Milford, MA, USA) ultraperformance LC (UPLC) system and an EASY-Spray PepMap C18 column (Thermo Fisher, ES800; 3-m bead size, 75 m by 150 mm) for reversed-phase chromatography. Peptides were eluted over a linear gradient over the course of 60 min from 2% to 50% (vol/vol) acetonitrile in 0.1% formic acid. MS and MS/MS spectra were acquired in a data-dependent mode with up to six higher-energy collisional dissociation (HCD) MS/MS spectra acquired for the most intense parent ions per MS. For data analysis, MS peak lists were generated with ProteoWizard MSConvert ^58^, and database searches were performed with Protein Prospector software, v.5.16.1 (http://prospector.ucsf.edu/prospector/mshome.htm, UCSF) ^35^ against the SwissProt human protein database (downloaded November 1, 2017). The database was concatenated with an equal number of fully randomized entries for estimation of the false-discovery rate (FDR). Database searching was carried out with tolerances of 20 ppm for parent ions and 0.8 Da for fragment ions. For database searches, peptides sequences were matched to tryptic peptides with up to two missed cleavages. Carbamidomethylation of cysteine residues was used as a constant modification and variable modifications included oxidation of methionine, N-terminal pyroglutamate from glutamine, N-terminal acetylation, and loss of N-terminal methionine. The following Protein Prospector score thresholds were selected to yield a maximum protein FDR of less than 1%: a minimum “protein score” of 15 and a minimum “peptide score” of 10 were used; maximum expectation values of 0.01 for protein and 0.05 for peptide matches were used. The list of proteins was further manually curated to include only proteins identified in at least 2 out of 3 biological replicates, and with a minimum of two unique peptides identified in at least one biological replicate. The t-test analysis was performed on the NSAF values for proteins, calculated as previously described ^36^: NSAF is the number of spectral counts (SpC, the total number of MS/MS spectra) identifying a protein, divided by the protein’s length (L), divided by the sum of SpC/L for all proteins in the experiment. GO pathway enrichment analysis was performed using the DAVID 6.8 database (https://david.ncifcrf.gov, accessed 01/2018) ^37^. Label-free quantitation was used to compare relative abundance of selected proteins. The MaxQuant software package ^59^ was used to obtain normalized peak areas for precursor ions from extracted ion chromatograms, using the MaxLFQ algorithm ^60^. Raw mass spectrometry data files and peak list files have been deposited with accession number MSV000082206 at ProteoSAFE (http://massive.ucsd.edu).

### Statistics and Data Availability

Statistical significance (p-values) was determined with paired Student’s *t*-test or one-way ANOVA for differences between experimental group mean values. Where data required non-parametric statistics, the Wilcoxon signed-rank test or the Mann-Whitney U test were used for paired or unpaired experimental groups (respectively). The statistical test used in each experiment is specified in the relevant figure legends. Survival curves were calculated according to the Kaplan–Meier method and compared using a univariate Cox– Mantel log-rank test. Cox regression multivariate analysis was used to determine if KHSRP expression was an independent predictor of survival (variables included: age, gender, size, stage), using SPSS version 20.0 (IBM, Armonk, New York, USA). Two-tailed p-values < 0.05 were considered statistically significant. Data are represented as mean ± S.E.M., unless stated otherwise. Analyses were performed using Prism 6 (Graphpad Software, La Jolla, CA, USA) or R Studio version 1.1.463 (RStudio, Inc., Boston, MA. http://www.rstudio.com/).

Gene expression microarray data from this study have been deposited in the Gene Expression Omnibus under accession number GSE112329. Raw data from the proteomic analysis is publicly available for download on the MassIVE database (ftp://massive.ucsd.edu/MSV000082206).

## RESULTS

### KHSRP is overexpressed in Colorectal cancer

To profile *KHSRP* expression in CRC, we performed a meta-analysis of CRC microarray datasets from the Oncomine database. *KHSRP* expression was increased in tumor compared to normal tissue across 18 CRC patient cohorts from 8 different datasets, with a median adjusted p-value of 4.06×10^-6^ (Figure 1-A, and supplementary table 1). *KHSRP* ranked in the top 5% of overexpressed genes in two of these analyses (from GSE9348 and GSE6988), and in the top 25% for 8 other analyses. In the remaining 8 analyses *KHSRP* was not one of the most overexpressed genes (<25%). Only 4 of the 18 analyses (from GSE5261 and GSE9689) fell below the threshold of significance. Figure 1-B shows individual data from the GSE9348 cohort^38^, in this dataset *KHSRP* expression was increased by 2-fold on average in 70 patients with early stage (stage II) colorectal carcinoma compared to healthy controls (p=4.6×10^-11^). In the TCGA cohort ^39^ (also shown in Figure 1-B) *KHSRP* expression increased in 211 samples of colorectal adenocarcinoma (any stage) compared to normal tissue (pooled analysis, +1.36 fold change, p<0.0001). In an additional analysis of the GSE6988 cohort ^40^, *KHSRP* expression increased in samples from either primary colon adenocarcinoma (n=25) or liver metastases (n=13) compared to matched normal colon mucosa and normal liver tissue, respectively (Figure 1-C).

**Figure 1:**
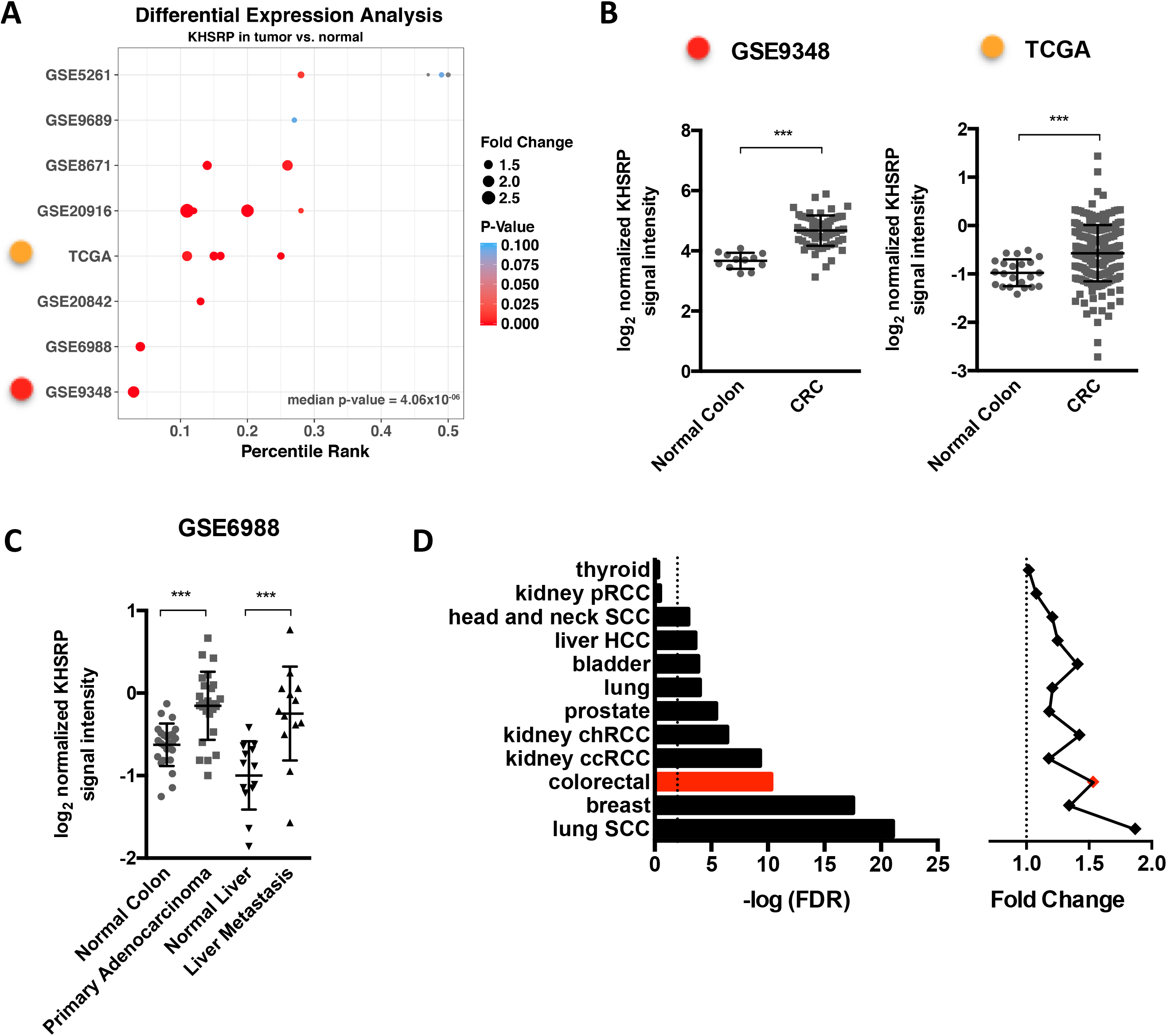
*In silico* analysis of publicly available CRC datasets reveals a consistent tumor-specific over-expression of KHSRP. **(A)** Differential expression analysis of KHSRP in the Oncomine database returns 18 analyses (each comprising a different patient group compared to normal tissue) across 8 different CRC datasets. The analyses are plotted by percentile ranking of KHSRP over-expression (as a percentage of all overexpressed genes in the dataset); dots are sized in proportion to the fold change of KHSRP expression in tumor vs. normal, and colored by p-value (further details of the analyses are provided in supplementary table 1). **(B)** Differential expression of KHSRP between tumor and normal tissue, showing individual data points along with the mean ± SD, is reported for the two representative analyses indicated (with the color code referring to panel A). *** p<0.0001 (Mann-Whitney test). **(C)** Differential expression of KHSRP between paired matched tumor and normal tissue in primary adenocarcinoma and liver metastasis. The mean ± SD is reported with individual points. *** p<0.0001 (Wilcoxon matched-pairs signed rank test). (**D**) Differential expression analysis for KHSRP in RNA-Seq data for the indicated 12 cancer types in the TCGA dataset. For each type, the fold change expression of KHSRP in tumor versus normal tissue and the associated FDR-adjusted p-value are reported

Increased *KHSRP* expression in tumor tissue was not limited to CRC but remained significant across a meta-analysis of 59 cancer types in the Oncomine database (median adjusted p-value=0.004, supplementary table 2). In addition, to look at expression of *KHSRP* across different cancer types we utilized RNA-Seq transcriptomic data available in the TCGA database, from which differentially expressed genes were derived for 12 different cancer types against their respective normal tissue ^23^: *KHSRP* was significantly overexpressed to varying degrees in 10 cancer types (Figure 1-D), with CRC being among the most highly significant (Fold change overexpression=1.53, FDR-adjusted p-value=5.02×10^-11^). The colon has also generally a high tissue-specific cancer expression of *KHSRP* across the entire spectrum of cancers represented in the TCGA (Supplementary Figure 1). However, the frequency of genetic alterations (mutations, deletions or amplifications) in *KHSRP* is less than 2% in the CRC TCGA dataset (Supplementary Figure 2), which is low considering that the three most commonly mutated genes in CRC (APC, TP53 and KRAS) have mutation frequencies of 81%, 60% and 43%, respectively ^39^. Collectively, these *in silico* data provided a framework to support additional investigations into the putative role of KHSRP in the pathogenesis of CRC.

### KHSRP is a marker of poor prognosis in colorectal cancer

Next, we investigated KHSRP protein expression in primary tissue samples. Protein extracts were obtained from fresh frozen colorectal tumor and normal adjacent tissue of 16 patients following surgical resection. WB analysis revealed a two-fold increased expression (p=0.0009) of KHSRP in stage IV CRC, compared to matched uninvolved (*adjacent normal*) tissue control (Figure 2-A). Analysis of stage II-III patients showed a similar trend (Supplementary Figure 3-A). The KHSRP antibody used for WB detected a single band at ∼82 kDa (Supplementary Figure 3-B), the predicted size for the protein, confirming the specificity of this antibody which has been previously orthogonally validated (Human Protein Atlas).

**Figure 2:**
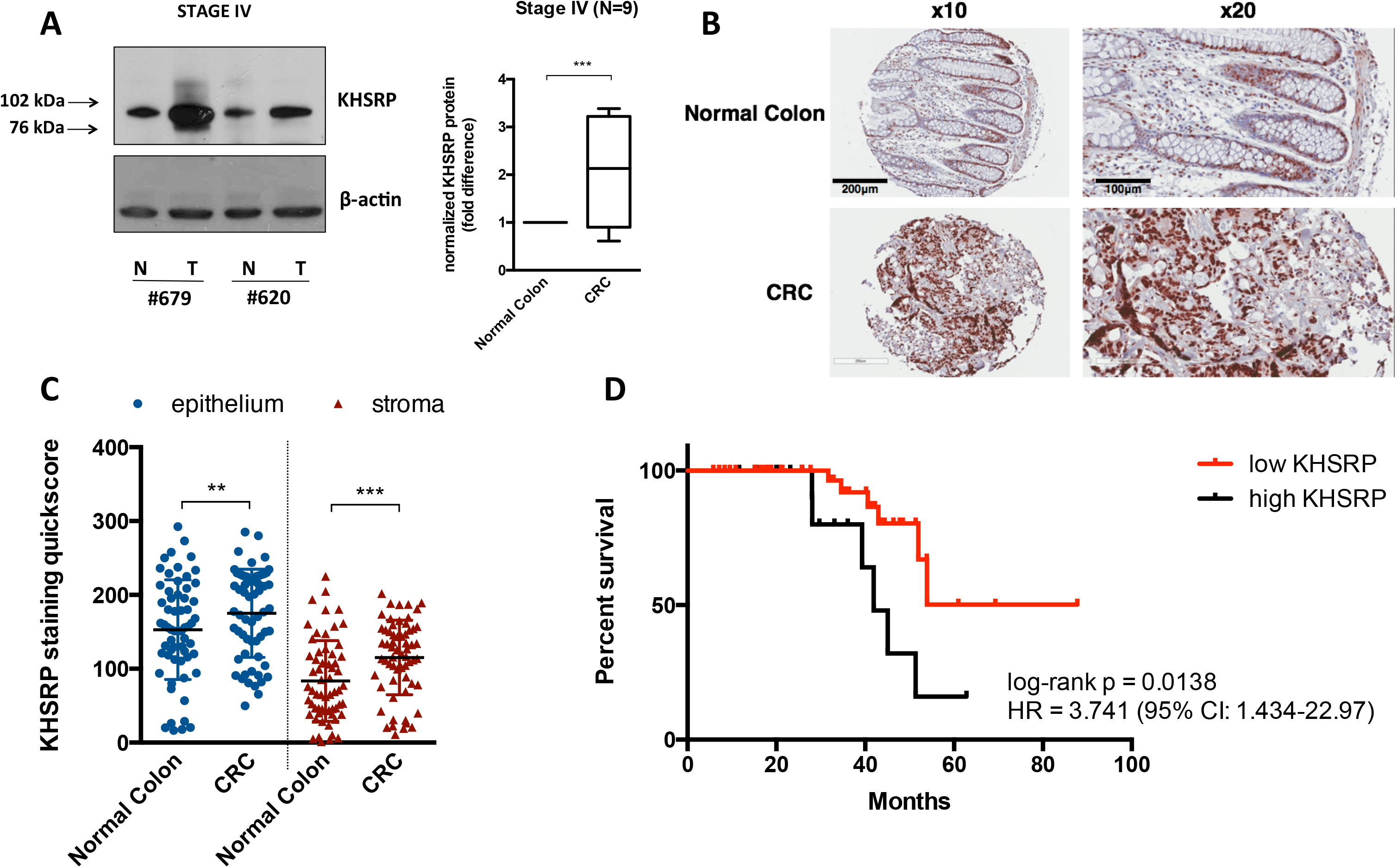
Analysis of KHSRP expression in primary tissue samples reveals a tumor-specific overexpression with prognostic value. **(A)** Western blot analysis of KHSRP in lysates from fresh-frozen tissue samples of patients with stage IV CRC (N=9), comparing tumor (T) and matched normal tissue (N) from each patient. A representative blot for 2 patients is shown along with semi-quantitative densitometric analysis of all patients (the Box and whisker plot depicts interquartile range, median, minimum and maximum limits) ***p<0.0001 (Student’s *t* test) **(B)** Representative images from TMA sections of normal and CRC tissue stained for KHSRP. **(C)** Quantitation of stromal and epithelial KHSRP staining in normal colon vs. primary CRC tissue samples from the TMA. The mean ± SD is reported with individual points. ** p<0.01 and ***p<0.0001 (Wilcoxon). **(D)** T/N ratios in the epithelium were used to segregate patients, using 20% as a cutoff (T/N<1.2 versus T/N>1.2). Survival curves were calculated according to the Kaplan–Meier method and compared using the Cox–Mantel log-rank test. The hazard ratio (HR) and associated confidence interval are shown.

To expand on these initial results, and to compensate for the limited number of samples analyzed by WB, we then performed immunohistochemistry (IHC) to investigate KHSRP expression in tissue obtained from a cohort of 62 patients with late stage (III-IV) CRC ^25^. KHSRP was expressed in both the epithelial and stromal compartments (Figure 2-B). KHSRP expression was significantly higher in tumor versus matched normal tissue in both epithelium and stroma (Figure 2-B and 2-C). This was mainly due to differences in percentage of positive cells, while the intensity values were similar overall (Supplementary Figure 3-C). There were more KHSRP positive cells in the epithelium than in the stroma (Figure 2-C), however the increased expression in tumor compared to normal was more prominent in the stroma, as indicated by a higher tumor-to-normal (T/N) ratio of KHSRP staining in the stroma (Supplementary Figure 3-D). When T/N ratios are plotted separately for each patient, the majority had increased KHSRP expression in the tumor compared to matched normal tissue (Supplementary Figure 3-C and D). High KHSRP expression (more than 20% increase in tumor vs. normal tissue) was associated with decreased overall survival (Figure 2-D, HR=3.74, 95% CI = 1.43-22.97, p=0.0138) in a univariate analysis. Multivariate Cox regression analysis demonstrated that higher KHSRP expression in tumor epithelium was an independent predictor of poor survival, associated with a 2.5-fold increase in disease-associated death (*p* = 0.009). As most of the patients in this cohort were diagnosed with metastatic disease, relapse-free survival was not included in the analysis. There was also a weak but significant correlation of the KHSRP T/N ratio with tumor size in both epithelium (Pearson r=0.43, p=0.0016) and stroma (Pearson r=0.38, p=0.0051).

For a limited number of cases (N=10 patients), tissue was also available for the corresponding liver metastasis. In both the epithelium and the stroma, the KHSRP score was significantly higher in the metastatic tumor tissue compared to matched normal liver (Supplementary Figure 4-A). Overall the metastasis-to-normal (M/N) ratio was positive for all except one patient, but the difference between metastasis and normal was vastly larger in the stroma than in the epithelium, with higher M/N ratios (Supplementary Figure 4-B, C, and D); this can be explained by the low expression of KHSRP in the normal liver stroma. Comparison of matched primary tumor and liver metastasis (n=10 patients) showed that the expression of KHSRP is unchanged in the epithelial cells upon migration, however the metastatic stroma has a higher expression compared to the stroma in the primary tumor (Supplementary Figure 4-E and F).

To validate the prognostic role of *KHSRP* in an independent cohort, we used a published dataset of 566 patients with stage I-IV CRC in the French *Cartes d’Identité des Tumeurs* program ^41^. *KHSRP* expression was increased in tumor compared to uninvolved (*normal*) tissue, in accordance with our previous data (Figure 3-A). Survival analysis was restricted to stage II-III patients following recommendations from the original authors: high *KHSRP* expression was associated with a decreased 5-year relapse-free survival in this cohort (Figure 3-B, HR=1.95, 95% CI = 1.21-2.60, p=0.0038). *KHSRP* expression was not associated with other tumor features such as loss of mismatch repair (MMR) proteins, or mutations in *KRAS* or *BRAF* (Figure 3-C).

**Figure 3:**
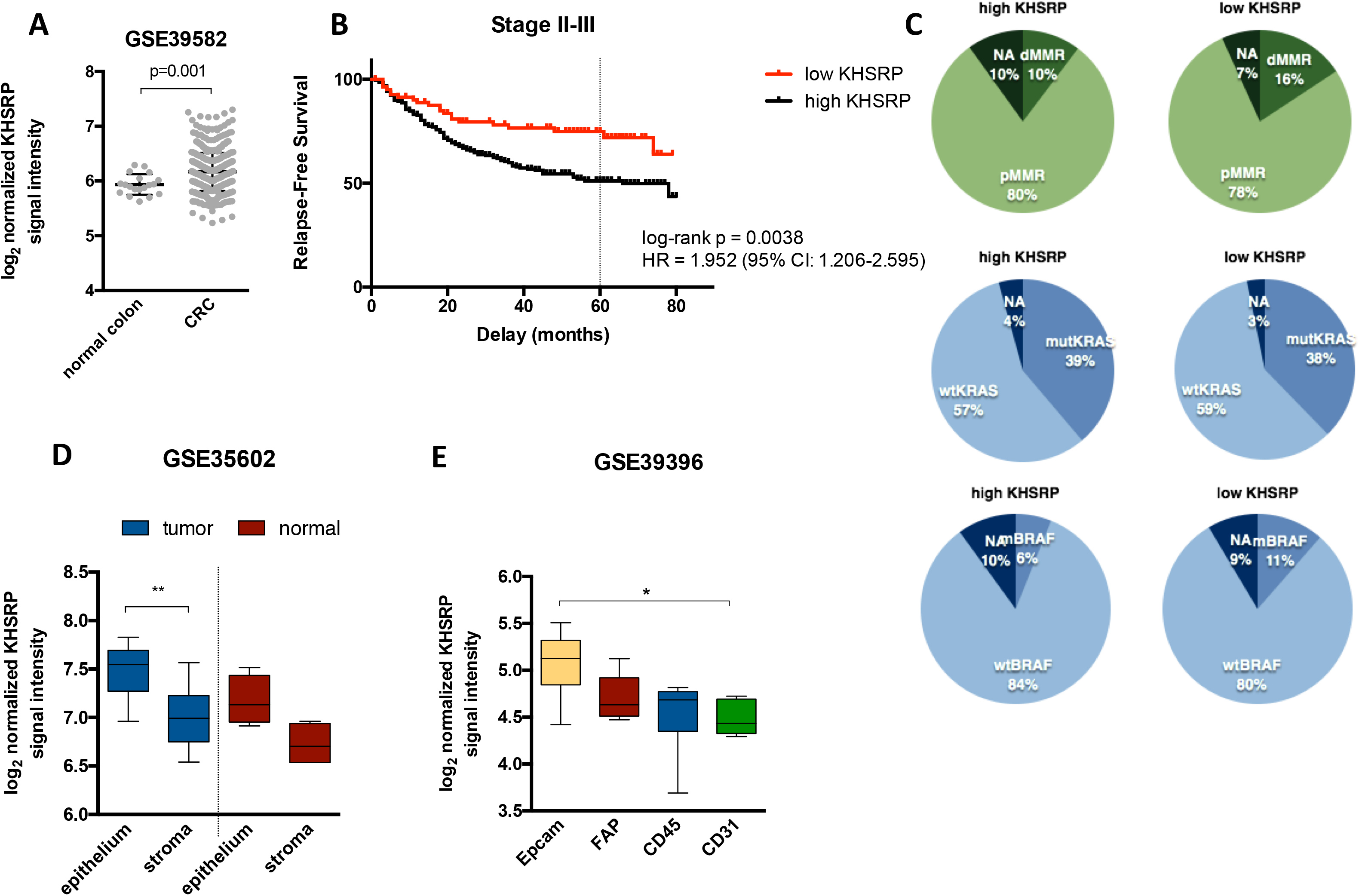
in silico analysis of three independent cohorts validates the overexpression and prognostic significance of KHSRP. **(A)** Log_2_ normalized signal intensity of *KHSRP* in a cohort of 566 CRC patients and 19 normal controls profiled by gene expression microarrays. The mean ± SD is reported with individual points and statistical significance (Mann-Whitney test). **(B)** Survival curves for patients segregated into high/low groups using the 25^th^ percentile of the *KHSRP* probe signal intensity. The hazard ratio (HR) and associated confidence interval are shown. **(C)** Distribution of patients with high/low *KHSRP* expression according to clinical characteristics: mismatch repair deficient (dMMR, i.e. MSI) or proficient (pMMR, i.e. MSS), KRAS (codon 12 and 13) mutational status, BRAF (p.V600E) mutational status. **(D)** Expression of *KHSRP* in laser-capture microdissected epithelial and stromal cells from 13 CRC and normal colonic mucosa samples. ** p<0.01 (Kruskal-Wallis) **(E)** Expression of *KHSRP* in a cohort of FACS-sorted cells from 14 CRC patients. EpCAM: epithelial cancer cells; CD45: leukocytes; CD31: endothelial cells; FAP: cancer-associated fibroblasts. *p<0.05 (Kruskal-Wallis). The Box and whisker plots in D and E depict interquartile range, median, minimum and maximum limits

To independently validate the expression of KHSRP in epithelial and stromal compartments from the IHC data, we took advantage of a transcriptomic dataset of 13 CRC and normal colonic mucosa samples, in which epithelial cells and stromal cells had been micro-dissected by laser capture and profiled separately ^42^. *KHSRP* was significantly overexpressed in the micro-dissected epithelial tumor compared with tumor stromal areas, and to a lesser extent in the epithelium compared with the stroma of the normal colonic mucosa, in accordance with our IHC data (Figure 3-D). Although there was a trend in increased *KHSRP* expression in tumor versus normal tissue for both the epithelium and the stroma, these comparisons were not statistically significant, probably due to the low number of samples. Finally, we used transcriptomic data from purified cell populations that were isolated from 14 dissociated human primary CRC samples ^43^. Fluorescence-activated cell sorting (FACS) was used to enrich for epithelial cancer cells (EpCAM^+^), leukocytes (CD45^+^), endothelial cells (CD31^+^) and cancer-associated fibroblasts (CAFs) (FAP^+^). The expression of *KHSRP* was higher in the Epcam^+^ epithelial cell population compared to the stromal cells, with the CD31^+^ endothelial cells showing the lowest *KHSRP* expression (Figure 3-E).

### KHSRP is involved in promoting the growth and survival of CRC cells

Next, we turned to an *in vitro* model of CRC to investigate the mechanistic role of KHSRP in epithelial cancer cells. We used the SW480 cell line, which was derived from a primary tumor of a moderately differentiated Duke B colon adenocarcinoma ^44^. These cells constitutively express KHSRP with a diffuse intracellular pattern (Supplementary Figure 5). An initial screen with a pool of 4 different siRNA sequences targeting *KHSRP* effectively silenced mRNA expression by 50% (Figure 4-A), and stably reduced proliferation when cells were monitored in real-time for 6 days (Figure 4-B). We then confirmed and extended these initial results using an independent siRNA that effectively knocked down KHSRP protein expression (Figure 4-C): silencing of KHSRP reduced cell growth as measured by neutral red uptake (Figure 4-D), reduced clonogenic potential measured by both number of colonies (surviving fraction) and total colony area (Figure 4-E), and reduced spheroid formation in 3D Matrigel cultures (Figure 4-F). Similar results were obtained by silencing KHSRP in an independent cell line model, SW620, which was isolated from a lymph node after multiple metastases occurred in the same patient (Supplementary Figure 6). Furthermore, we also developed a conditional (doxycycline-dependent) shRNA-based stable knockdown model from both SW480 and SW620 cell lines, which further confirmed results obtained with the transient silencing model (Supplementary Figure 7).

**Figure 4:**
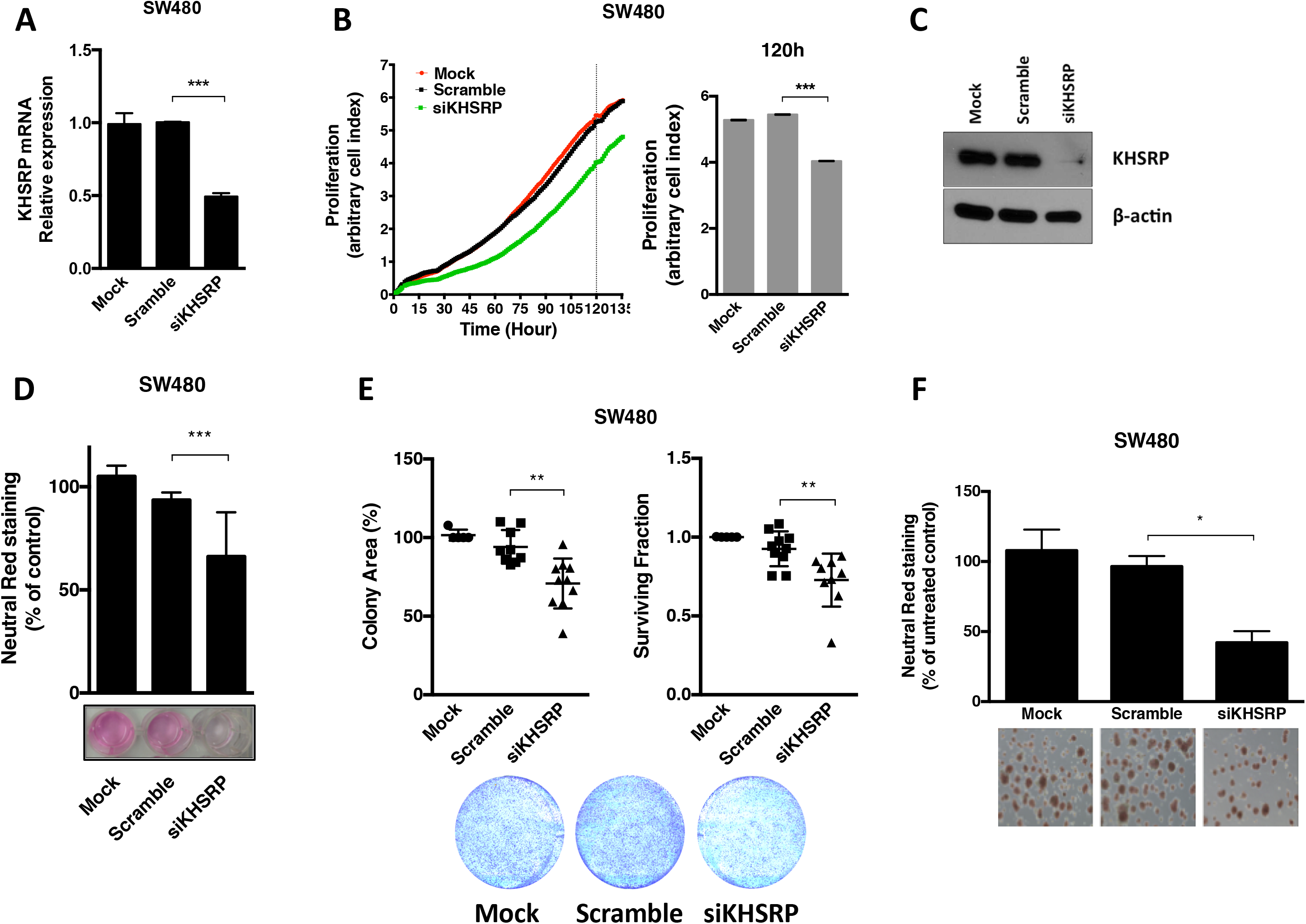
KHSRP is involved in growth and survival of an *in vitro* CRC cell line model. **(A)** qRT-PCR analysis of SW480 cells transfected with a pool of siRNAs targeting *KHSRP* (siKHSRP), a scramble control pool (Scramble), or transfection reagent control (Mock) for 48h. **(B)** Cells were transfected as in A, and continuously monitored for proliferation for the indicated time; full time-course growth traces are shown, along with quantification of cell index (arbitrary measure of cell proliferation based on impedance measurements) at the indicated time point. **(C)** WB analysis of KHSRP protein expression in SW480 cells transfected with an siRNA targeting *KHSRP*, or a scramble control siRNA, or a mock (transfection reagent) control for 48h. **(D)** Growth of cells transfected as in C, monitored after 7 days by neutral red assay; representative images from one experiment are shown, with staining quantification from triplicate assays. Growth is expressed as % of untreated control (cells with no transfection) **(E)** Clonogenic potential for cells transfected as in C, measured by both colony area and surviving fraction. representative images from one experiment are shown, with quantification from triplicate assays. **(F)** Spheroids from cells transfected as in C, grown in Matrigel; representative images are shown with staining quantification from triplicate assays. Growth is expressed as % of untreated control (cells with no transfection) (**A-F**)**p<0.001, and ***p<0.0001 (ANOVA) for all tests in the figure.

### KHSRP regulates cell cycle and affects the tumor-promoting microenvironment

To investigate the mechanisms involved in the tumor-promoting effects of KHSRP, we started by analyzing the transcriptomic profiles of siRNA-transfected KHSRP knockdown SW480 cells, compared to scramble control transfection (Supplementary Figure 8). This preliminary analysis identified 135 genes differentially regulated upon KHSRP knockdown (Supplementary Table 3), which are enriched for functional categories related to cell growth and survival (specifically, cell cycle and apoptosis), as well as cell signaling and known KHSRP-related functions (translation, protein metabolism), as shown by GO terms analysis (Supplementary Figure 8-B). Three representative genes (*FABP3, MTF1, SMG1*) were further validated by qRT-PCR (Supplementary Figure 8-C). The protein interactions inferred from the differentially expressed transcripts (Supplementary Figure 8-D) ^31^ included gene networks for cell cycle, transcription and protein synthesis, further suggesting a role for KHSRP in cell cycle control. This prompted us to investigate the effects of KHSRP knockdown on the cell cycle distribution of SW480 cells. KHSRP siRNA caused an increase in the G0/G1 population (Figure 5-A), concomitant with a decrease in the mitotic index, measured as the number of cells positively stained by phospho-Histone H3 (Figure 5-B), indicative of a reduced proliferation rate due to a delayed G1/S transition. These data, together with that presented in Figure 4, suggest an involvement of KHSRP in driving epithelial cells into mitosis, thereby increasing proliferation. Furthermore, stable knockdown of KHSRP resulted in reduced expression of c-Myc (Supplementary Figure 7-C).

**Figure 5:**
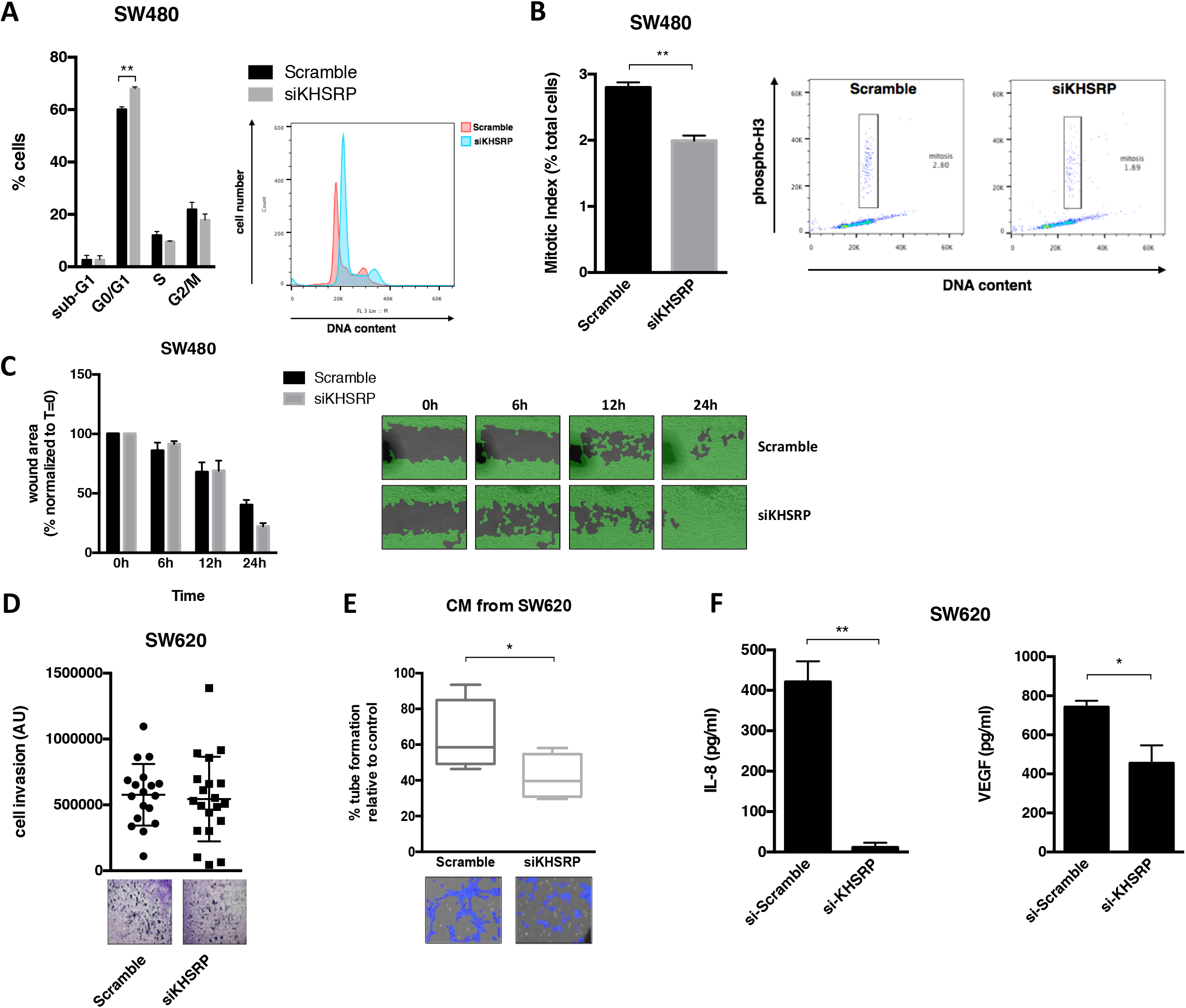
KHSRP is involved in regulating cell cycle and mitosis, as well as influencing the pro-tumorigenic extracellular environment *in vitro*. **(A)** Cell cycle analysis of SW480 cells transfected with siKHSRP or Scramble negative control siRNA; a representative histogram of propidium iodide staining from one experiment is shown, along with a bar graph of cell distribution into cell cycle phases across three replicate experiments. **p<0.001 (ANOVA) **(B)** Analysis of mitotic cells by quantitation of phosphor-histone H3 in SW480 cells transfected as in A; a representative histogram is shown with bar graph of three replicate experiments. **p<0.001 (Mann-Whitney). **(C)** Wound-healing assay of SW480 cells transfected as in A; representative images from one experiment are shown, with quantitation of wound area from two replicate experiments. **(D)** Invasion of SW620 cells transfected with siKHSRP or Scramble negative control siRNA, measured by the capacity to migrate through Matrigel; quantification of invaded cells from three replicate experiments is shown, along with representative images. **(E)** Endothelial tube formation of HDEC cells treated with conditioned media (CM) from SW620 cells transfected as in D. Tube formation is expressed as % of control (HDEC cells treated with the same media without previous conditioning in SW620 cells) *p<0.05 (Student’s *t* test). **(F)** Expression of secreted IL-8 and VEGF in the CM of SW620 cells transfected as in D, measured by ELISA. *p<0.05, **p<0.001 (Student’s *t* test).

KHSRP has been involved in regulating cell migration and invasion in different non-colorectal tumor models ^72^. However, silencing of *KHSRP* did not influence cell migration (as measured by a wound-healing assay) in SW480 cells (Figure 5-C), and did not impact on the ability of the metastatic SW620 cells to invade through a Matrigel layer (Figure 5-D). Nevertheless, the cell culture supernatant from SW620 cells with silenced KHSRP was able to reduce endothelial cell growth in a tube-forming assay (Figure 5-E), suggesting that KHSRP may regulate the secretion of specific extracellular signaling mediators promoting angiogenesis. In support of this, CRC cells with silenced KHSRP expression secreted reduced levels of IL-8 and VEGF (Figure 5-F).

### KHSRP regulates the secretion of multiple proteins by tumor cells

To further investigate the involvement of KHSRP in regulating the extracellular microenvironment, we employed the conditional KHSRP knockdown cell line model (described in Supplementary Figure 7) to probe the secretome of the metastatic SW620 cell line upon doxycycline-induced, shRNA-mediated knockdown of KHSRP. Shotgun proteomics analysis identified 191 proteins, of which 27 where unique for the control samples (i.e. undetected in the doxycycline-treated samples), and 16 where unique for the doxycycline-treated samples (undetected in the control samples). Statistical analysis of spectral counts data showed that 40 proteins were significantly under-expressed or over-expressed by more than 50% upon doxycycline treatment; the majority of these differentially regulated proteins (33 out of 40) were down-regulated (Figure 6-A, and Supplementary Table 4). GO term analysis of the 40 highly differentially regulated proteins revealed an enrichment for known terms related to KHSRP function (e.g. RNA binding, spliceosome) and expected terms related to the cellular location of the probed sample (e.g. membrane and extracellular space); interestingly, there was also a significant enrichment for terms related to the pro-tumorigenic extracellular microenvironment (e.g. exosomes, focal adhesion, cell migration), as well as terms broadly related to the immune response and specifically to myeloid cell activation (Figure 6-B). Using label-free quantitation, we validated specific examples of candidate proteins whose secretion is deregulated upon KHSRP knockdown (Figure 6-C). Downregulated proteins included: the Rho-specific guanine nucleotide dissociation inhibitor-α (RhoGDIα), a master regulator of Rho GTPases that promotes cell migration ^45^; the S100 calcium binding protein A11 (S100-A11), an endoplasmic reticulum-associated calcium-binding protein that regulates cell death ^61^; Ephrin-B2, a membrane-anchored proteins which activates paracrine tyrosine kinase signaling impacting proliferation, migration and response to therapy ^62^. Conversely, upregulated proteins included: the actin-binding protein Villin-1, a protein which upon cellular stress induces intestinal epithelial cell death by necroptosis, as well as increased inflammation ^46^; ADAMTS-like protein 2, an extracellular matrix-residing protein with roles in matrix remodeling and modulation of metalloprotease activity ^63^. The KHSRP-dependent downregulation of protein S100-AA11 and Ephrin-B2 was further confirmed by Western blot (Figure 6-D).

**Figure 6:**
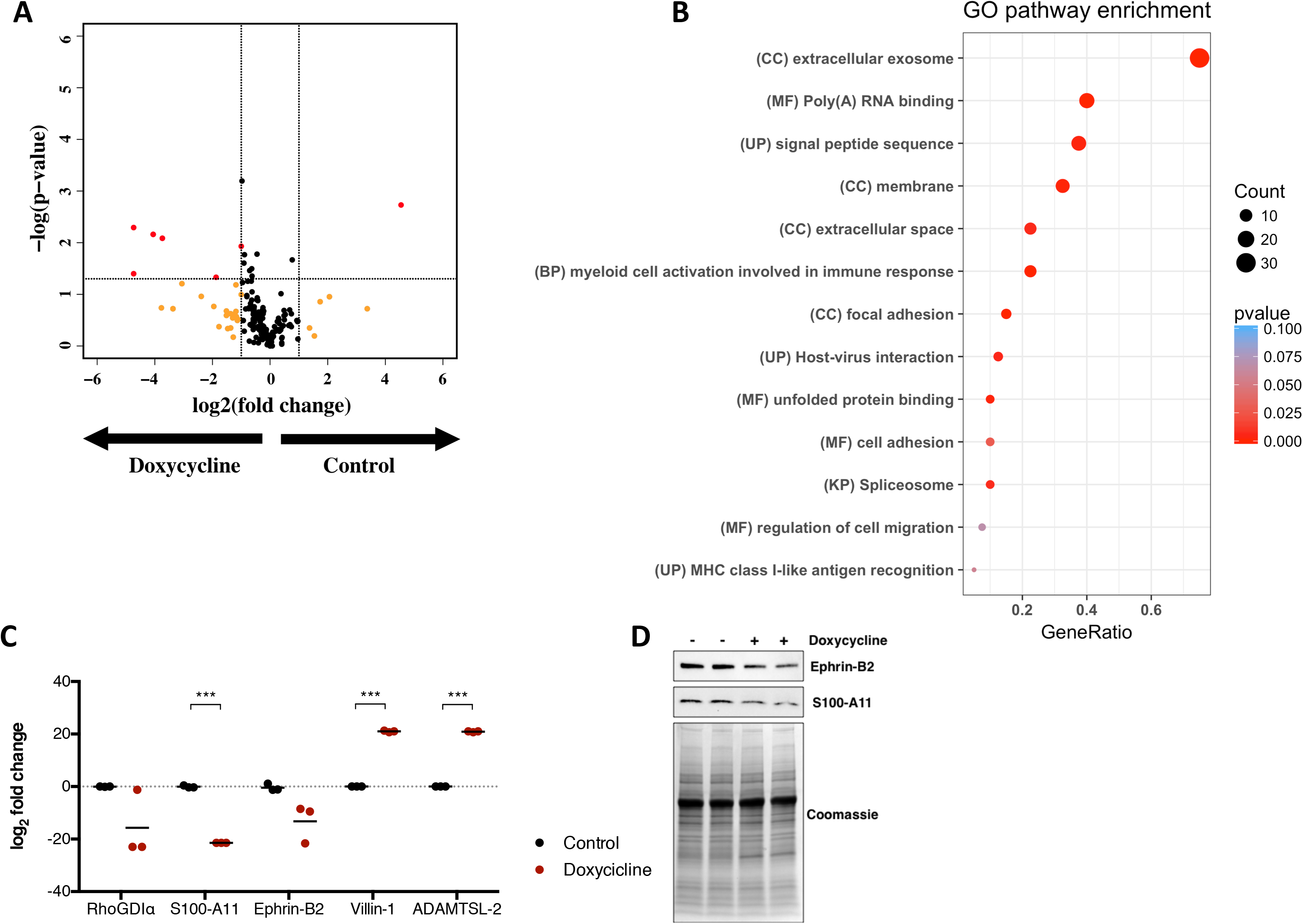
KHSRP regulates expression of key proteins that modulate the tumorigenic microenvironment. Differential expression of secreted proteins was analyzed by shotgun proteomics in conditioned media from SW620 cells treated with Doxycycline for 4 days to induce shRNA-mediated KHSRP knockdown. **(A)** Volcano plot showing differentially regulated proteins in the doxycycline-treated samples compared to control, colored in red (log_2_ Fold Change > |1|, p-value < 0.05). Proteins that are differentially regulated above the fold change threshold with a non-significant p-value are colored in yellow **(B)** Dot plot of results from a GO analysis of the 40 highly differentially regulated proteins. GO terms are reported with their category (CC=cellular component, MF=molecular function, BP=biological process, UP=uniprot annotation, KP=Kegg pathway) and plotted by gene ratio (the number of genes in one GO term compared to the total). Dots are sized in proportion to the number of hits within a GO term, and colored by p-value. **(C)** Label-free quantitation of relative abundance of the indicated five proteins in the doxycycline-treated vs. control samples. ***p<0.0001 (Student’s *t* test). **(D)** Western blot analysis of Ephrin B2 and protein S100-A11 in conditioned media from control or doxycycline-treated SW620 cells. A Coomassie stain is shown for protein loading control.

## DISCUSSION

The role of KHSRP in the pathogenesis and progression of CRC has not been extensively studied so far. Here, we provide orthogonal evidence supporting a pro-tumorigenic role of KHSRP in CRC, based on extensive mining of multiple publicly available data sets, analysis of patient samples and *in vitro* cell line models. Our results are in agreement with recent reports in osteosarcoma ^13^, lung cancer ^12, 47^, papillary thyroid cancer ^67^, and esophageal squamous cell carcinoma ^48^.

Our IHC data provide evidence for KHSRP overexpression in the context of both primary tumor and metastasis. However, we surprisingly found that KHSRP remained unchanged in the metastatic epithelial cells infiltrating the liver compared to that of the primary site, while stromal expression was strongly upregulated from a very low basal expression in the normal hepatic stroma. This suggests that invading epithelial cells are capable of modifying the local microenvironment, or alternatively that a local increase in KHSRP expression pre-exists and might contribute to the metastatic niche ^49^. In fact, KHSRP is involved in the transient expression of the chemokine CX3CL1 in liver epithelial cells in response to IFN-γ stimulation ^50^, contributing to the control of cellular homeostasis during inflammation and macrophage infiltration. Our proteomic data provide additional evidence in support of a putative role for KHSRP in regulating the tumor microenvironment, with implications for tumor progression and metastasis. To the best of our knowledge, this represents the first report of such an effect for KHSRP in shaping the extracellular environment of epithelial cells. Although preliminarily, our data also suggest a possible role for KHSRP in regulating the immune response, both at the gene-expression level and at the secretome level (as reported here by microarray and proteomic data). This suggests the intriguing idea that the reported negative prognostic role of KHSRP could be partly ascribed to a regulation of immune evasion mechanisms, and provides a rationale for reconciling the oncogenic roles of KHSRP with its other known functions in shaping innate immunity during viral infection and inflammation. In fact, recent work from Bezzi and colleagues using genetically engineered mouse models of prostate cancer driven by the loss of specific tumor suppressor genes has shown that the genetic landscape of epithelial tumor cells can directly influence the composition of immune cell infiltrates in primary tumors, through mechanisms based on distinct chemokine pools resulting from aberrant transcriptional and signaling programs ^51^. Furthermore, knockout mice studies have shown a requirement of KHSRP for the post-transcriptional control of type-1 and 3 IFN genes ^52, 68^. Recent mechanistic evidence proved that KHSRP can directly associate and negatively regulate retinoic acid-inducible gene I (RIG-I) receptor signaling during viral infection; this exert a negative control on the activation of type-I IFN innate immune responses during the recognition of pathogen-associated molecular patterns (PAMPs) encoded by viral RNA ^53^. This could have potential consequences for the control of anti-tumor innate immune responses, as similar mechanisms are involved in the recognition of damage-associated molecular pattern (DAMPs) and the immunogenicity of tumor cells ^54^. Consistently, KHSRP has been implicated in the innate immune response to oncogenic pathogens such as *H. pylori* in the stomach ^69^ and HPV in the uterus ^70^.

One question that remains open is how overexpression of KHSRP is controlled, given that our data suggest it is not the product of a driver mutation event. A possible explanation could come from studying another RBP which is commonly overexpressed in CRC, Musashi (MSI)-1 ^64^. Spears and colleagues have demonstrated a reciprocal interaction between MSI-1 and APC through a double-negative feedback loop, whereby MSI-1 contributes to APC inactivation, an early initiating driver event in the majority of sporadic CRC ^65^. This suggests the intriguing hypothesis that increased expression of RBPs (perhaps not limited to MSI-1) might co-evolve as a passenger non-mutational event early in the adenoma-to-carcinoma sequence. Another hypothesis, related to the effect of KHSRP on the tumor microenvironment that our data begins to suggest, is that deregulation of RBPs could be selected for indirectly through negative regulation of immune surveillance, a relevant force shaping cancer evolution. In support of an early event hypothesis, our *in silico* data suggest that KHSRP expression is unchanged in hypermutated, microsatellite-instable CRC, a type of tumor characterized by a proportionally increased burden of passenger mutations ^66^. Nevertheless, specific mutations in KHSRP could still impact on protein functions and show increased frequencies in some cancer types ^72^. Furthermore, although limited to a very small fraction of the dataset, the analysis from the Oncomine database provides information on colorectal mucinous carcinoma, a distinct form of CRC found in 10-15% of patients and characterized by distinct molecular features and mutation events ^71^ in this subset of patients, *KHSRP* was not one of the most overexpressed genes, suggesting that it might be an event specifically related to adenocarcinoma.

There is also considerable interest in the suitability of different members of the translational control machinery as potential therapeutic targets in cancer, although the evidence so far is at the preclinical stage ^55^. Small molecule inhibition of the RBP HuR in APC^Min^ mice led to a reduction in small intestinal tumor formation, and a concomitant reduction in c-myc expression ^56^. Another recent report showed that inhibition of mTORC1 using rapamycin caused regression of established APC-deficient intestinal tumors, suggesting that inhibition of translation elongation using existing clinically approved drugs might benefit patients at high risk of developing CRC ^57^. Our loss-of-function data in *in vitro* CRC models suggest that KHSRP could also have putative therapeutic implications and warrants further investigation into the suitability of such an approach.

In conclusion, our report sheds light onto the molecular role of KHSRP in CRC, providing for the first time comprehensive data in support of a pro-tumorigenic role of this RBP through direct modulation of epithelial cell phenotype and indirect modulation of the tumor microenvironment.

## ADDITIONAL INFORMATION

### Ethics Approval and Consent to Participate

All human tissue used in this study was obtained with the informed written consent of the patient; ethical approval was granted by the Research and Ethics Committee of St. Vincent’s University Hospital (Dublin, Ireland). The study was performed in accordance with the Declaration of Helsinki.

### Authors contribution

Study concept and design: FC, EJR, JOS, GAD; acquisition of data: FC, KO, MT, LB, FON, RP, SN, PM; analysis and interpretation of data: FC, KO, JP, KK, SF, KS, JOS, EJR; drafting and critical revision of the manuscript: FC, KO, JOS, EJR, CSC; funding and support: EJR, KS, GAD, DF, CSC; administrative, technical or material support: MT, BN; study supervision: EJR, GAD.

## Acknowledgments

We gratefully acknowledge Dr. Giselle Knudsen (UCSF), Dr. Sudipto Das (RCSI), Dr. Janet McCormack (UCD), Dr. Barry Moran (TCD), and Dr. Dimitri Scholz (UCD) for technical assistance at various stages of this project. We also wish to thank Dr. Martin Kampmann (UCSF) for providing advice and material for the generation of the conditional knockdown cell lines, and Dr. Quing Xiong (Southwest University, Chongqing, China) for providing access to their data. The bioinformatic expertise of Dr. Tim Hacker and Sam Ivry (UCSF) is also acknowledged. Mass spectrometry was performed in collaboration with the UCSF Mass Spectrometry Facility (directed by Prof. Alma Burlingame).

## Notes

**Financial Support:** This work was supported by a Newman Fellowship awarded by the UCD Foundation to FC funded by a donation from Merck Serono, from a Science Foundation Ireland Industry Fellowship (14/IFB/2715) to FC, NIH P41CA196276 to CSC and from the Irish Health Research Board (HRA _POR_2013_281) to EJR. The sponsors had no role in the study design, collection, analysis, or interpretation of data.

#### Summary of Updates

This version of the manuscript reflects revisions that occured during multiple peer review rounds from June 2018 to June 2019.

ftp://massive.ucsd.edu/MSV000082206

https://www.ncbi.nlm.nih.gov/geo/query/acc.cgi?acc=GSE112329

