## Supplementary Figures and Tables for "KH-type splicing regulatory protein controls colorectal cancer cell growth and modulates the tumor microenvironment"

**SUPPLEMENTARY INFORMATION**

**Supplementary Table 1:** details of the differential expression analysis for KHSRP in CRC datasets from the Oncomine database. For each analysis the corresponding dataset and citation is reported, along with the KHSRP gene rank, p-value (Student’s *t*-test, corrected for multiple hypothesis testing using the false discovery rate method), fold change, and percentile rank.

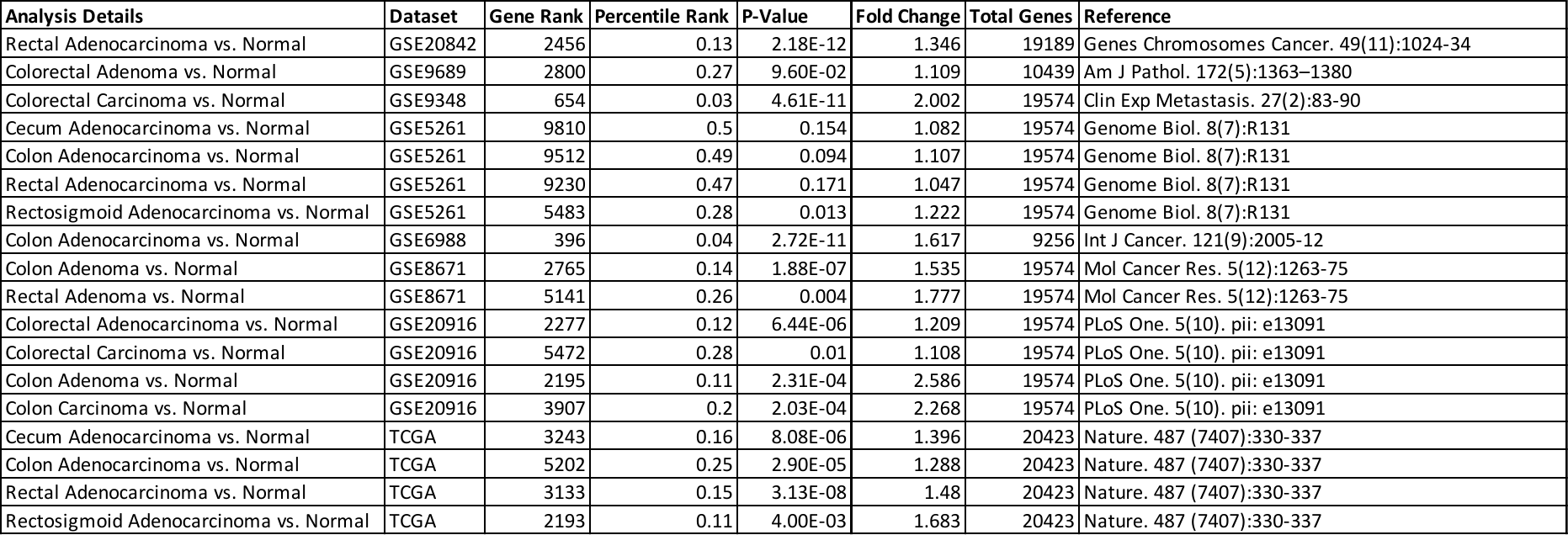

**Supplementary Table 2 (next page):** details of the differential expression analysis for KHSRP in pan-cancer datasets from the Oncomine database. Each analysis is reported with the associated dataset in Oncomine, the corresponding citation, and the associated percentile rank.

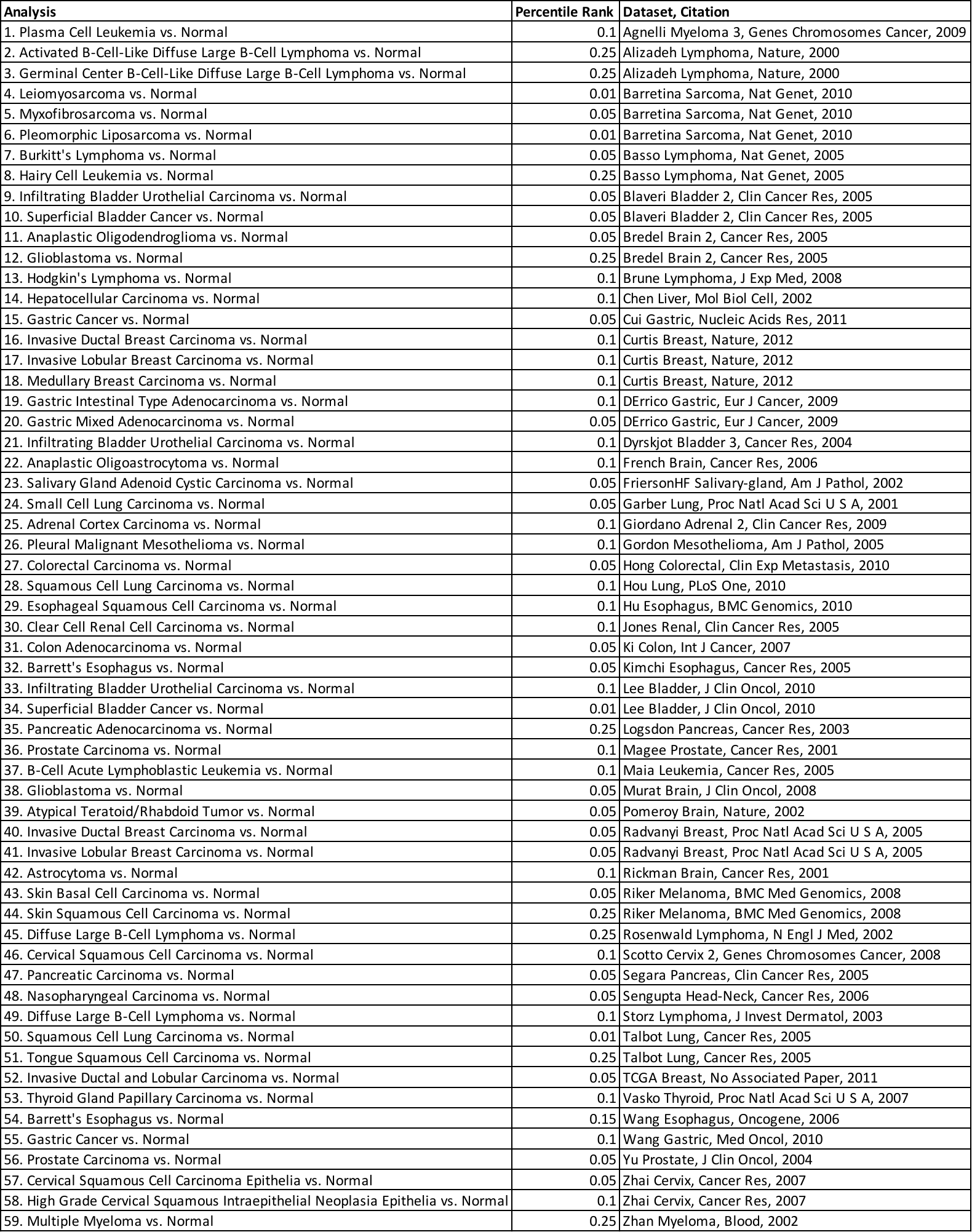

**Supplementary Table 3 (continues on the next page):** list of the 135 differentially regulated genes in the microarray. Log_2_-normalized fold change, and associated p-values, are reported for the comparison of SW480 cells transfected with siKHSRP versus siSCR control.

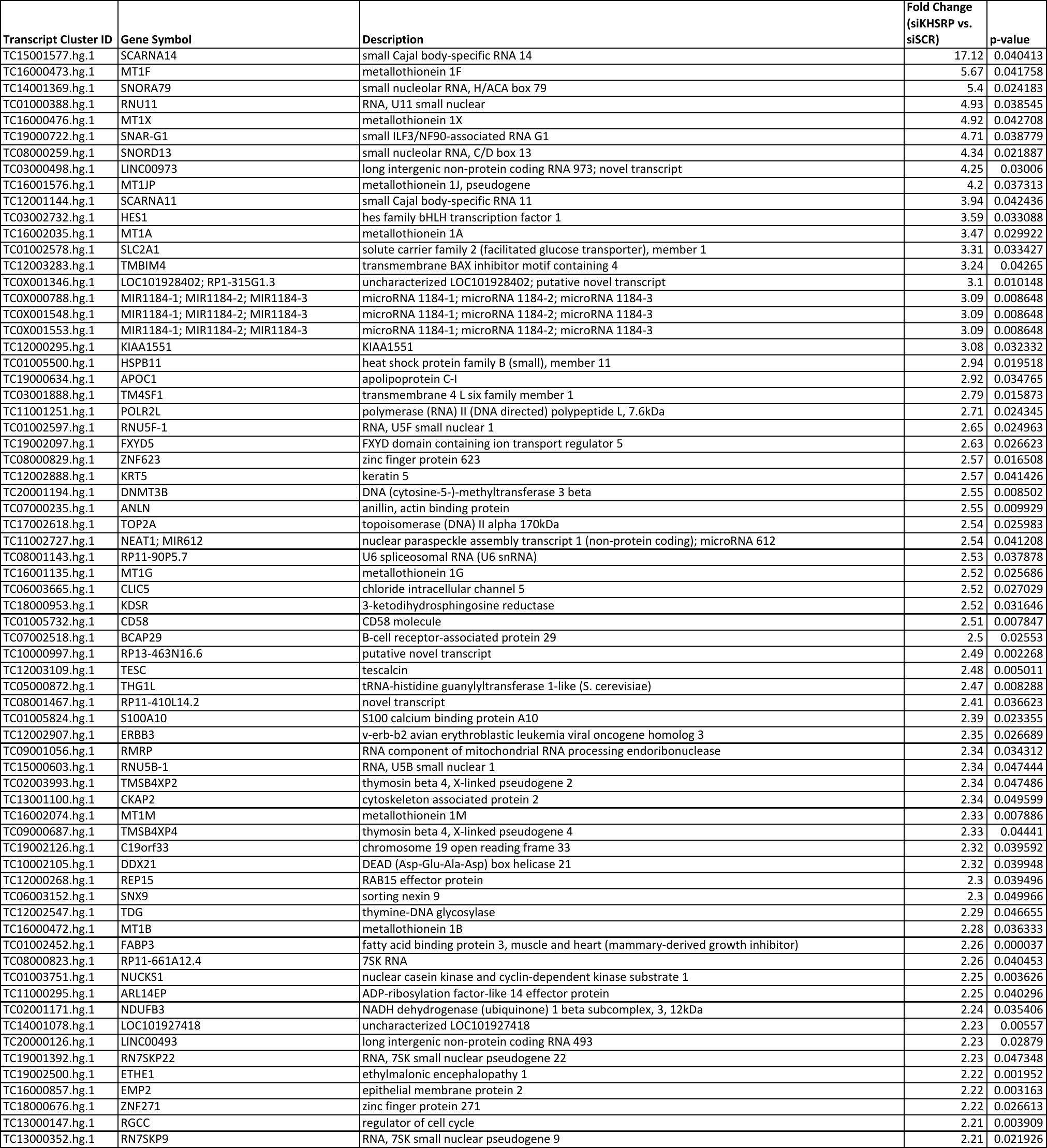

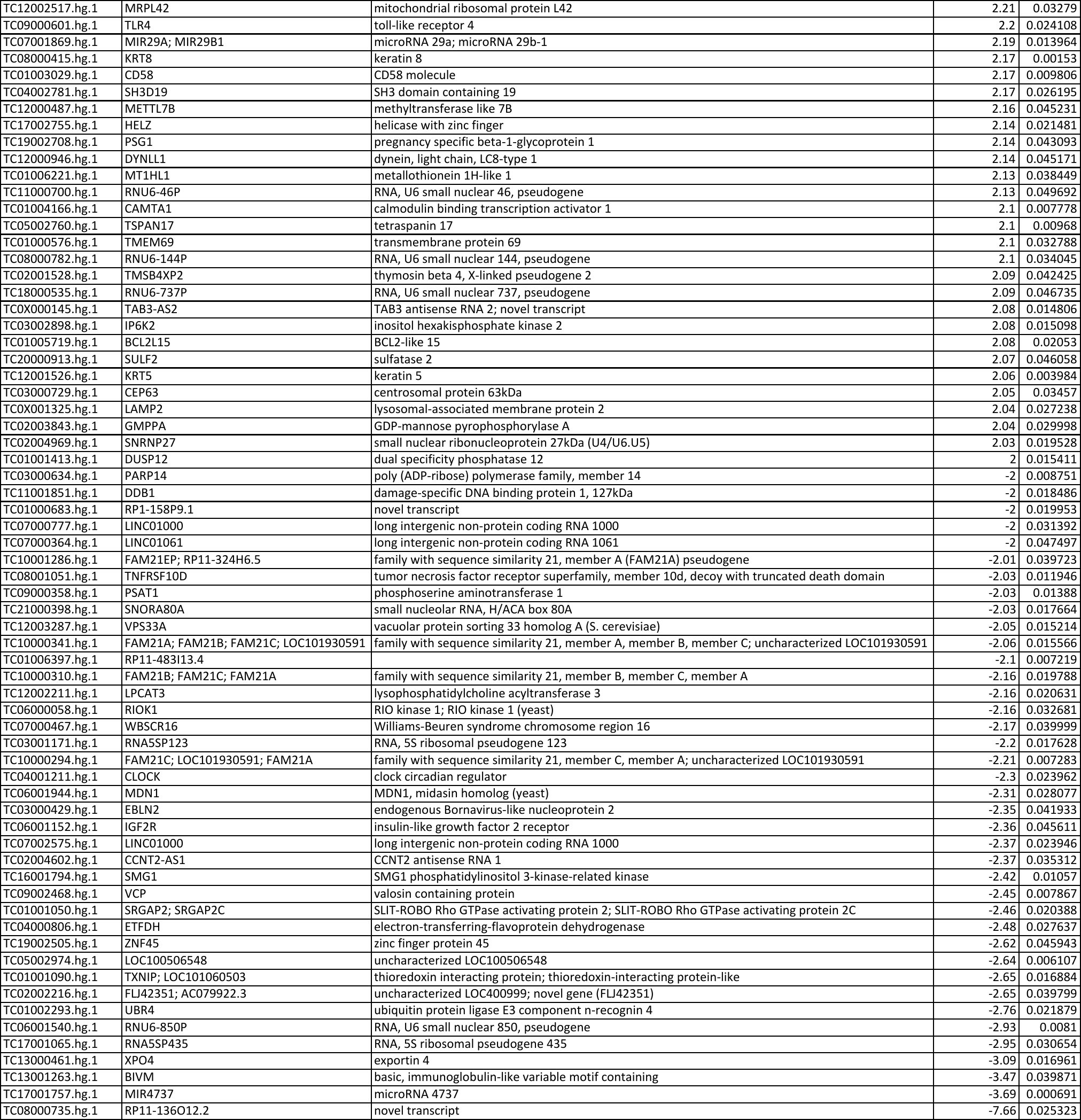

**Supplementary Table 4:** list of the 40 differentially regulated proteins in the shotgun proteomics experiment. Log_2_-normalized fold change, and associated p-values, are reported for the comparison of doxycycline- versus vehicle (ctrl)-treated SW620 cells. GenBank accession numbers (ACCNUM) and Entrez Gene Identifiers (ENTREZID) are also reported.

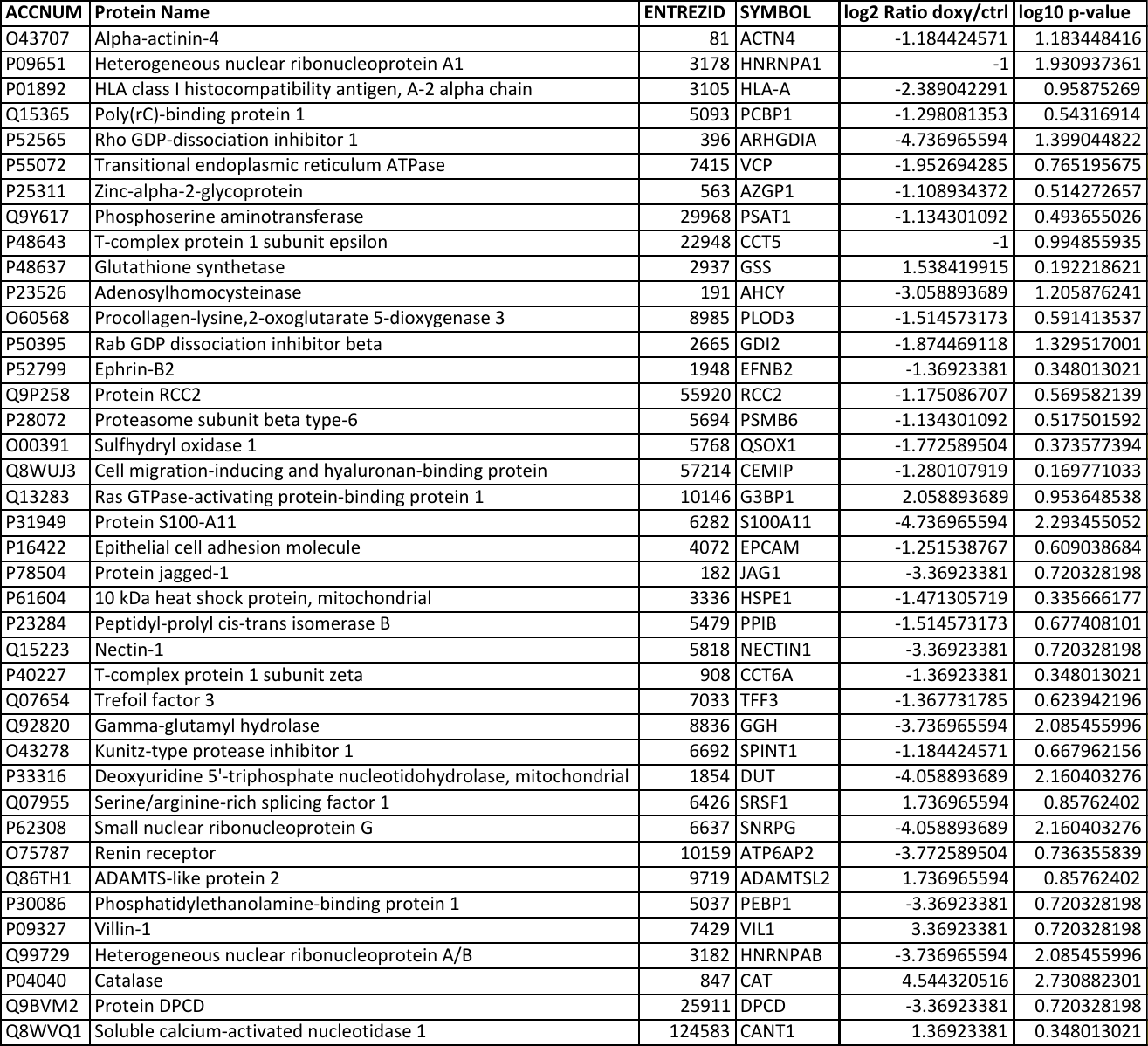

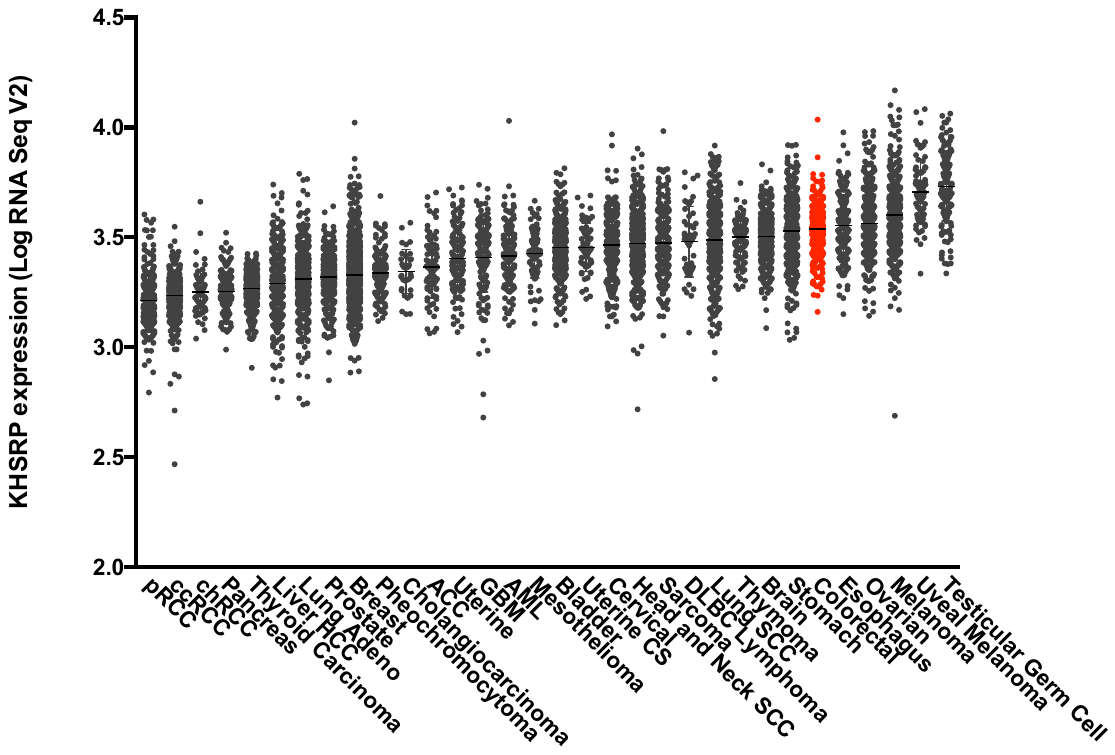

**Supplementary Figure 1:** RNA-Seq expression data for KHSRP in the TCGA cohort was plotted in increasing order, grouped by tumor type (for details of the tumor type codes refer to the TCGA Data Portal, https://tcga-data.nci.nih.gov/docs/publications/tcga/)

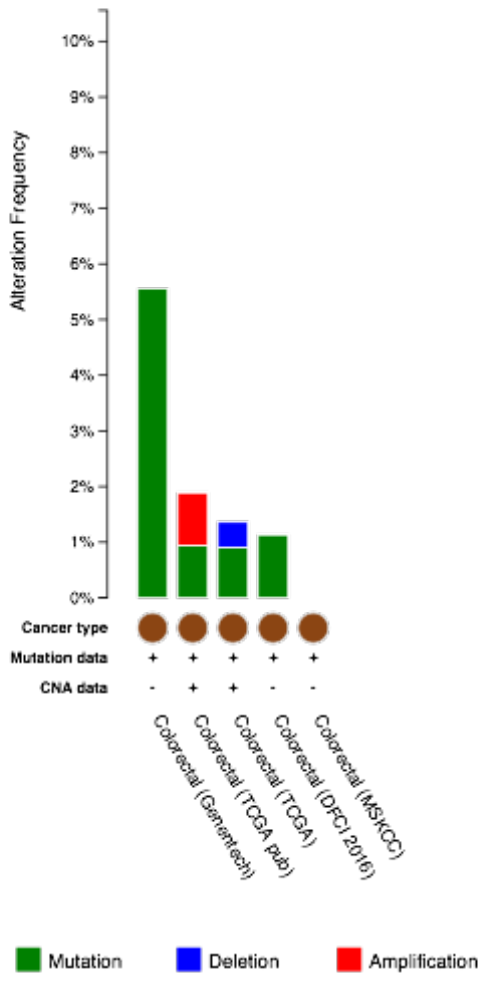

**Supplementary Figure 2:** The frequency of genetic alterations in KHSRP across 4 different CRC datasets as reported in the cBio portal: DFC|2016 (dbGaP accession: phs000722), Genentech (European Genome-Phenome Archive accession: EGAS00001000288), MSKCC (dbGaP accession: phs000790.v1.p1), and TCGA.

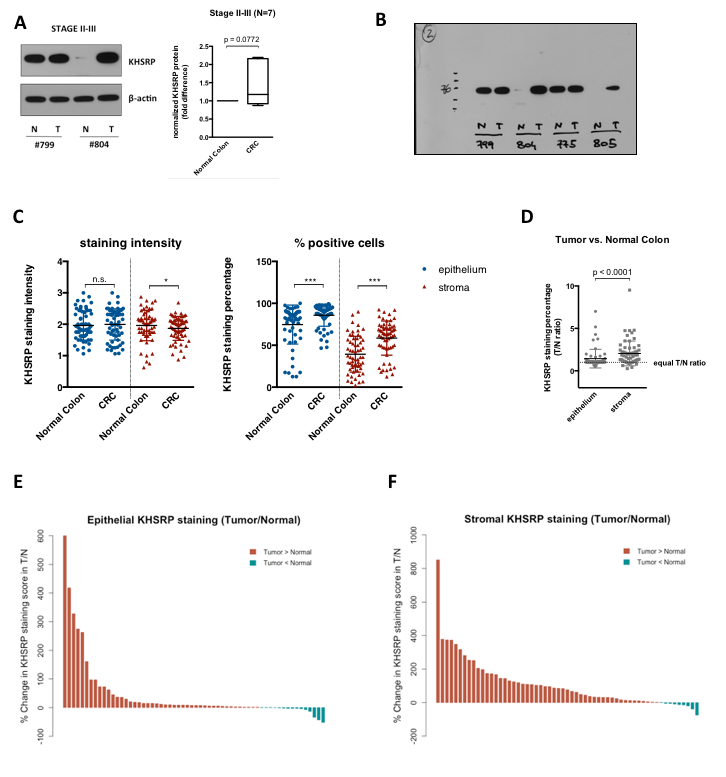

**Supplementary Figure 3: (A)** Western blot analysis of KHSRP in lysates from fresh-frozen tissue samples of patients with stage II-III CRC (N=7), comparing tumor (T) and matched normal tissue (N) from each patient. A representative blot for 2 patients is shown along with semi-quantitative densitometric analysis of all patients. The p value (Student’s *t* test) is indicated. (**B**) Full-size image of the blot for KHSRP corresponding to panel A. (**C**) Separate quantification of staining intensity and percentage between normal and tumor tissue, for both epithelium and stroma. **(D)** Quantification of tumor-to-normal (T/N) ratio of KHSRP staining in epithelial and stromal compartments of TMA. Waterfall plots of T/N ratios in the epithelium **(E)** or stroma **(F)** indicate patients with increased or decreased tumor-specific KHSRP expression.

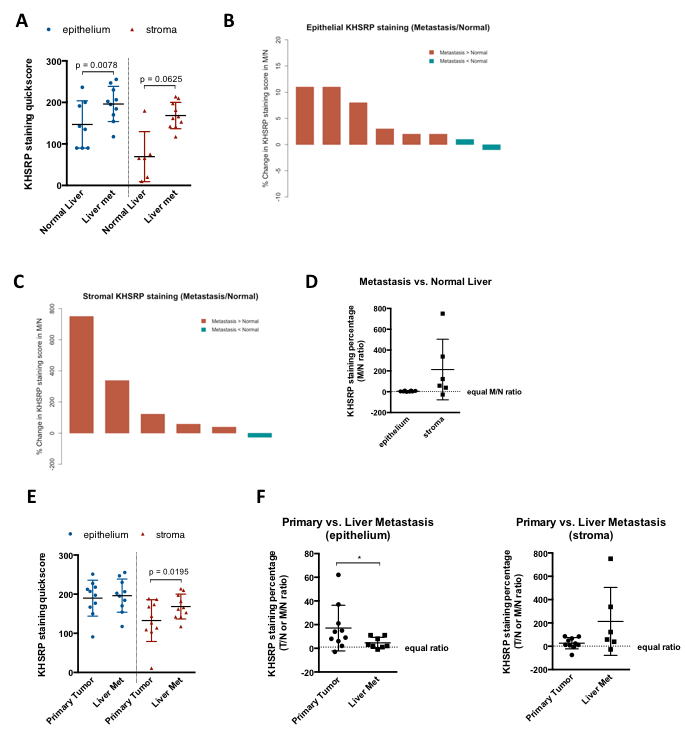

**Supplementary Figure 4: (A)** Quantitation of stromal and epithelial KHSRP staining in normal liver vs. liver metastasis tissue from CRC patients from the TMA. **(B-C)** Quantification of metastasis-to-normal (M/N) ratio of KHSRP staining in epithelial and stromal compartments of TMA. Waterfall plots of M/N ratios for single patients are shown, along with **(D)** a comprehensive plot of all patients (N=8). **(E)** Quantitation of stromal and epithelial KHSRP staining in primary CRC tumor vs. liver metastasis tissue for 10 patients from the TMA. **(F)** Comparison of T/N ratios in the primary tumor with the M/N ratios in the corresponding liver metastasis for both epithelial and stromal compartments of 10 patients in the TMA.

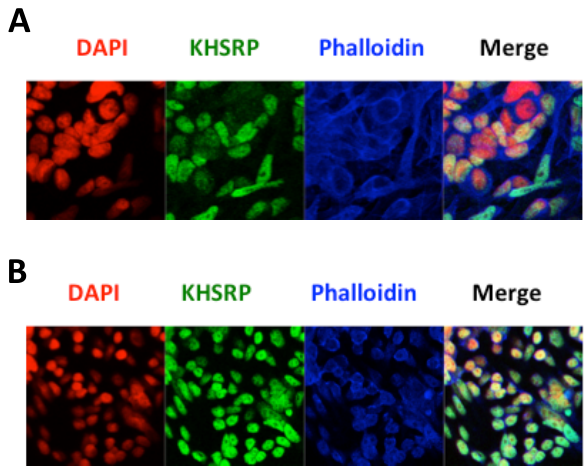

**Supplementary Figure 5:** Immunofluorescent images of **(A)** SW480 and **(B)** SW620 cells. Red: nuclei (DAPI), Green: KHSRP, Blue: F-actin (Phalloidin).

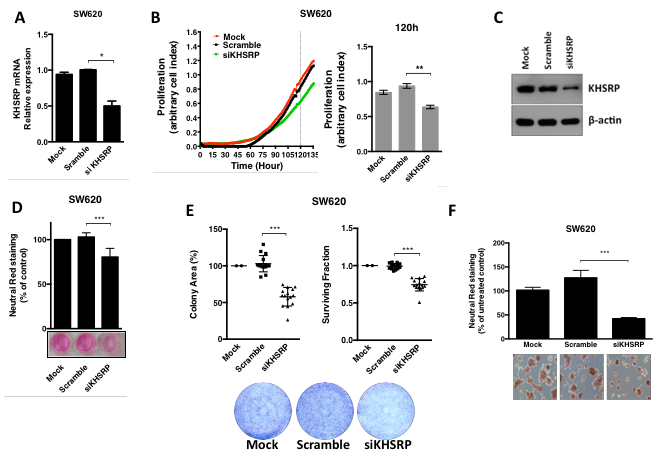

**Supplementary Figure 6: (A)** qRT-PCR analysis of SW620 cells transfected with a pool of siRNAs targeting *KHSRP* (siKHSRP), a scramble control pool (Scramble), or transfection reagent control (Mock) for 48h. **(B)** Cells were transfected as in A, and continuously monitored for proliferation for the indicated time; full time-course growth traces are shown, along with quantification of cell index (arbitrary measure of cell proliferation based on impedance measurements) values after 5 days. **(C)** WB analysis of KHSRP protein expression in SW620 cells transfected with an siRNA targeting *KHSRP*, or a scramble control siRNA, or a mock (transfection reagent) control for 48h. **(D)** Growth of cells transfected as in C, monitored after 7 days by neutral red assay; representative images from one experiment are shown, with staining quantification from triplicate assays. **(E)** Clonogenic potential for cells transfected as in C, measured by both colony area and surviving fraction. representative images from one experiment are shown, with quantification from triplicate assays. **(F)** Spheroids from cells transfected as in C, grown in Matrigel; representative images are shown with staining quantification from triplicate assays. (**A-F**)*p<0.05, **p<0.001, and ***p<0.0001 (ANOVA) for all tests in the figure.

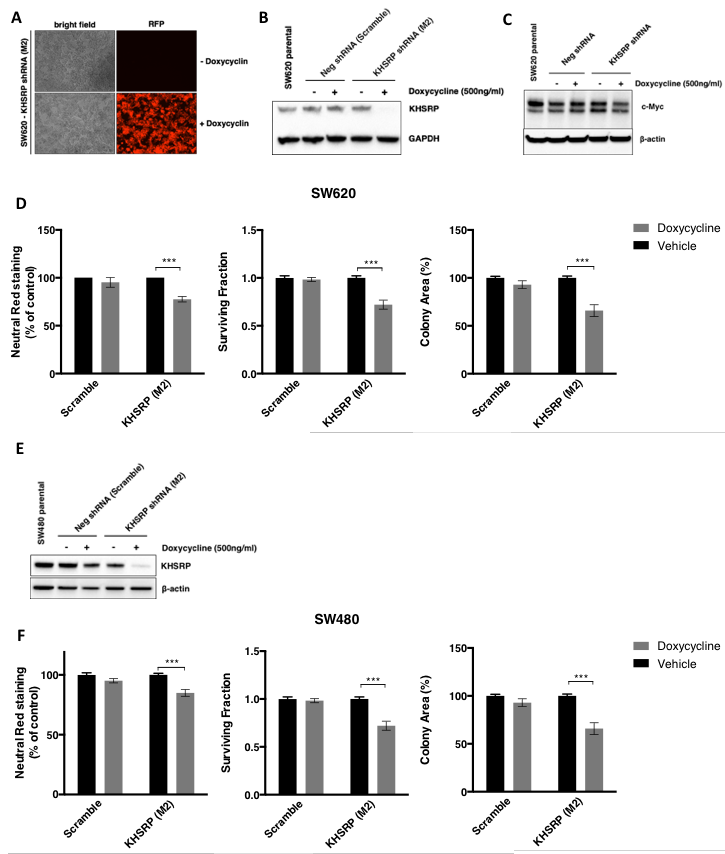

**Supplementary Figure 7:** SW620 and SW480 cells stably transfected with a conditionally-expressible shRNA pool targeting KHSRP (M2) or a non-targeting control pool (Scramble). **(A)** Phase-contrast and fluorescence images of SW620 cells treated with doxycycline or vehicle control for 4 days. Consistent expression of red fluorescent protein (RFP) indicates induced expression of the shRNA-containing cassette, and **(B)** WB analysis confirmed effective knock down of KHSRP protein expression. **(C)** WB analysis of the same cells revealed reduction of c-Myc protein expression upon knock down of KHSRP. **(D)** Growth of stably transfected SW620 cells in the presence and absence of doxycycline was quantified using the neutral red assay; colony formation was measured by both surviving fraction and colony area; ***p<0.0001 (ANOVA). **(E)** WB analysis confirmed effective knock down of KHSRP protein expression in SW480 generated and treated as above. **(F)** Growth of stably transfected SW480 cells in the presence and absence of doxycycline was quantified using the neutral red assay; colony formation was measured by both surviving fraction and colony area); ***p<0.0001 (ANOVA).

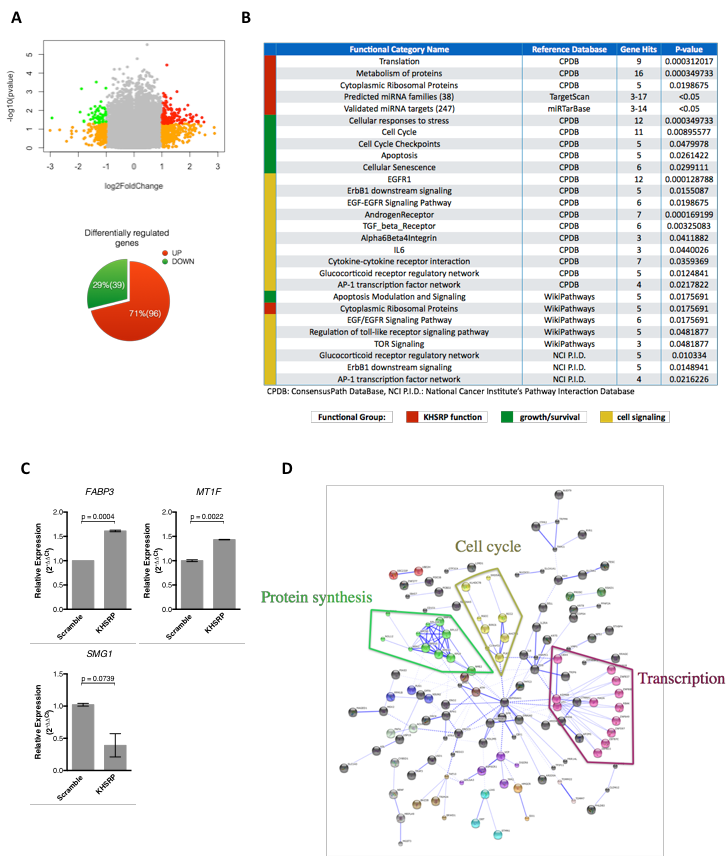

**Supplementary Figure 8 (previous page):** transcriptomic profile of SW480 cells transfected with KHSRP siRNA compared to scramble negative siRNA control. **(A)** Volcano plot showing differentially regulated genes in green (down-regulated) and red (up-regulated) (log_2_ Fold Change > |1|, p-value < 0.05). Genes that are differentially regulated above the fold change threshold with a non-significant p-value are colored in orange. The distribution of the 135 significantly differentially regulated genes is depicted in the pie chart, using the same color scheme. **(B)** Over-representation analysis of the 135 differentially regulated genes. GO terms are arbitrarily grouped by biological function, and reported with the reference database, the corresponding number of gene hits and the associated p-value. (**C**) qRT-PCR analysis of the indicated genes in SW480 cells stably transfected with a conditionally-expressible shRNA pool targeting KHSRP or a non-targeting control pool (Scramble), stimulated with doxycycline for 4 days to induce shRNA expression. The p value (Student’s *t* test) is indicated in each panel. **(D)** Network map of predicted associations for the protein products of the 135 differentially regulated genes. Proteins are represented by nodes (smaller nodes indicate proteins with unknown structural information); edges represent the predicted functional associations, with the thickness of the line indicating the degree of confidence for the prediction of the interaction. Proteins are color-coded arbitrarily by clustering, with the three main clusters annotated.
